# Dissecting the Dynamic Evolution of Tensional Homeostasis in Fibroblasts using an Integrated Biomechanical Bioreactor Platform

**DOI:** 10.64898/2026.02.23.707598

**Authors:** Andrew V. Glick, Victor V. Nguyen, Daniel Paukner, Margherita De Marzio, Huocong Huang, Girgis Obaid, Christian J. Cyron, Jacopo Ferruzzi

**Affiliations:** Department of Bioengineering University of Texas at Dallas, Richardson, TX, USA; Institute for Continuum and Material Mechanics, Hamburg University of Technology, Hamburg, Germany; Channing Division of Network Medicine, Brigham and Women’s Hospital and Harvard Medical School, Boston, MA, USA; Department of Surgery, University of Texas Southwestern Medical Center, Dallas, TX, USA; Department of Immunology, University of Texas Southwestern Medical Center, Dallas, TX, USA; Department of Biomedical Engineering, University of Texas Southwestern Medical Center, Dallas, TX, USA

**Keywords:** tensional homeostasis, bioreactor, mechanobiology, fibroblast, collagen remodeling

## Abstract

Mechanical homeostasis indicates the remarkable ability displayed by cells in tissues to maintain their mechanical properties near a stable homeostatic set-point. Experimental investigations and theoretical studies indicate that mechanical stress represents a key homeostatic target that stromal cells, such as fibroblasts, seek to maintain by tuning the intracellular structure and by remodeling the extracellular matrix. Much of what is known about mechanical homeostasis of tissues under tension, or tensional homeostasis, is based on experiments on tissue equivalents, that is fibroblast-populated collagen gels. However, existing platforms used to investigate tensional homeostasis cannot infer mechanical stress dynamically. Here we developed an integrated biomechanical bioreactor combining force sensing with confocal and multiphoton microscopy to dissect the mechanobiological mechanisms of tensional homeostasis. We used our platform to test the hypothesis that fibroblasts maintain a constant state of stress across varying collagen concentrations. Contrary to this assumption, synchronized force and imaging measurements revealed that stress is not constant but rather elevated at low collagen concentrations, where fibroblast contraction drives earlier collagen alignment and greater tissue compaction.

Conversely, force generation and α-SMA expression increase with increasing collagen concentration, accompanied by modest transcriptional changes. However, at the highest collagen concentration, this homeostatic balance is disrupted, with lower force generation and α-SMA expression, as gene expression shifts toward VEGFC-mediated autocrine survival signaling. These findings demonstrate that tensional homeostasis emerges from a dynamic balance between cellular contractility and extracellular matrix densification rather than stress maintenance while revealing that excessive matrix density disrupts this balance by triggering a pro-survival response.

## 1. INTRODUCTION

Tissue homeostasis is an essential requirement for multicellular life [1] and requires control over cell proliferation, differentiation, biochemical signaling, and physical interactions with other cells and with the surrounding extracellular matrix (ECM). A finely tuned control over these factors is key for maintaining local tissue variables to a nearly constant, or homeostatic, set-point [2]. Conversely, breakdown of homeostatic control leads to loss of function and disease across multiple tissues and organs [3]. Among the many physiological variables under homeostatic control, the mechanical state of connective tissues is maintained to operate under a preferred level of mechanical tension [4]. Fibroblasts represent the primary cell type in connective tissues responsible for maintaining mechanical homeostasis around a tissue-specific set-point [5,6]. Such a biomechanical set-point is thought to be established during embryonic development, maintained through cell-ECM interactions and eventually disrupted by maladaptive tissue remodeling. During disease progression, resident fibroblasts excessively produce and reorganize a collagen-rich ECM, leading to tissue stiffening and chronic fibrosis in vital organs [7]. Understanding mechanical homeostasis is therefore critical for improving antifibrotic treatments. However, it remains unclear how interactions between fibroblasts and a complex network of extracellular collagen give rise to a mechanically homeostatic state compatible with normal tissue function and how such a state is disrupted during fibrosis.

Much of what is known about mechanical homeostasis of tissues under tension, or tensional homeostasis, is based on experiments on tissue equivalents, that is fibroblast-populated collagen gels [8]. Free-floating tissue equivalents were originally developed as a model of wound contraction [9] and shown to compact the collagen matrix through contractile forces. While these constructs remain traction-free, fibroblasts can contract a circular gel to a fraction of its original diameter in the attempt to maintain mechanical homeostasis [10]. While doing so, fibroblasts remodel collagen heterogeneously and generate a residually stressed biomaterial [11]. For this reason, tissue equivalents anchored to a rigid substrate were introduced to balance the active stress generated by contracting fibroblasts [5]. Anchored tissue equivalents undergo macroscopic height reduction [12] while at the microscale, cell-generated traction forces reorganize collagen fibrils, generating a complex state of mechanical tension within the construct. This mechanical tension can be released by detaching the tissue equivalents, which triggers rapid changes in cytoskeletal organization and DNA synthesis [13] and leads to increasing rates of apoptosis in the absence of mechanical stress [14,15]. These early studies highlighted the need to quantify mechanical stress as fibroblasts remodel the surrounding collagen network. To this end, tissue equivalents restrained at one end to a rigid boundary and at the other to a force transducer were shown to develop tensile forces under static conditions [16], suggesting the existence of a homeostatic tensile force [17]. However, it is conceivable that the homeostatic target is a material quantity, such as the true (Cauchy) stress, rather than force alone. To gain a mechanistic understanding of how tensional homeostasis is developed and maintained in healthy soft tissues, and subsequently disrupted by disease processes, there is a need for improved tissue engineering platforms. Recent studies have presented engineered bioreactors capable of culturing fibroblasts in 3D collagen matrices under both uniaxial and biaxial tension [18]. However, current bioreactor designs have limited imaging capabilities, restricting structural measurements to discrete endpoints [19] rather than enabling continuous monitoring of both dynamic force generation and microstructural remodeling.

To address current limitations, we developed a miniaturized, computer-controlled biomechanical bioreactor fully integrated with confocal microscopy for real-time mechanobiological analysis. Our platform enables dynamic measurements of force evolution, local collagen remodeling, and global tissue compaction, thereby allowing dynamic stress estimation in tissue equivalents. In this study, we first validated the bioreactor’s ability to quantify mechanical stress and then used it to test the hypothesis that fibroblasts achieve tensional homeostasis by maintaining a constant state of stress in different collagen microenvironments. Contrary to the original hypothesis, our integrated analysis of mechanical stress evolution, collagen remodeling, contractile marker expression, and transcriptional profiling, demonstrates that fibroblasts do not maintain constant stress across collagen concentrations. Instead, tensional homeostasis emerges from a concentration-dependent balance between cell contractility and matrix compaction. When collagen concentration is too high, the proposed homeostatic balance is disrupted by reduced proliferation, decreased contractility, and the establishment of an autocrine signaling loop that promotes cell survival. Thus, this work establishes both an experimental platform and a framework for investigating the origins and disruption of tensional homeostasis.

## 2. MATERIALS AND METHODS

### 2.1 Biomechanical bioreactor

The bioreactor platform presented here is based on the design originally described by Eichinger et al. [18]. Major updates in actuation, force sensing, environmental control, and data acquisition were introduced to integrate synchronous biomechanical and imaging measurements. A schematic and photos of the device operating within a confocal microscope (Nikon, AX-SHS) are provided in Figure 1. Key components and features of the device are described below and in the supplementary methods.

**Figure 1.**
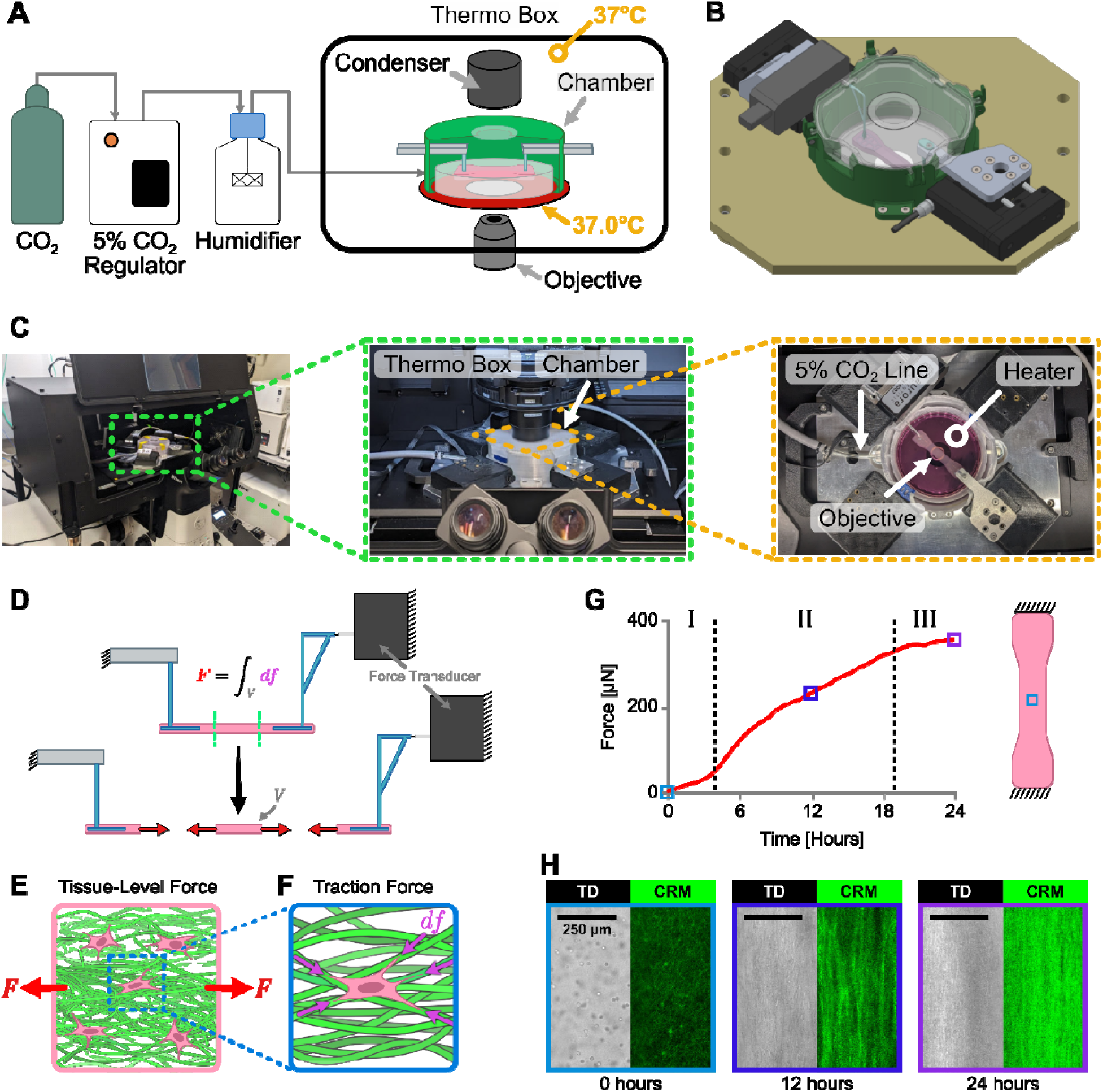
A biomechanical bioreactor fully integrated with a confocal microscope. (A) Schematic of the novel bioreactor platform integrated inside a confocal microscope. The unit is capable of sterile tissue culture, biomechanical sensing, and microstructural imaging of tissue equivalents for up to 72 hours. (**B**) SolidWorks rendering of the complete bioreactor assembly. (**C**) The bioreactor mounted inside a confocal microscope with key components labeled. **(D)** A free body diagram of the central region of a uniaxially constrained gel illustrates how **(E)** the tissue-level force (*F*) measured by the force transducer arises from **(F)** individual traction forces (*df*) exerted by fibroblasts at cell-ECM contacts. (**G**) Representative tissue-level force generated by fibroblasts within a uniaxially constrained collagen gel (1.0 mg/mL, 500,000 cells/mL) held at fixed length over 24 hours with **(H)** accompanying confocal reflection microscopy (CRM) and transmitted light detector (TD) images. Time-lapse confocal imaging synchronized with force recording demonstrates progressive collagen densification and fiber alignment along the axis of the bioreactor.

### 2.1.1 Biocompatible parts

Bioreactor components in direct contact with biological materials were 3D printed using a biocompatible resin (FormLabs, FLBMCL01) using a stereolithography (SLA) 3D printer (FormLabs, Form 4B). All parts were printed and washed according to manufacturer’s instructions. Briefly, after 3D printing parts were washed in a bag filled with 99% isopropyl alcohol (IPA) in an ultrasonic bath for 15 minutes, flipping the bag halfway through. The parts were then soaked in fresh 99% IPA for 5 minutes, and sonicated for 2 minutes in fresh 99% IPA, flipping the bag halfway through. Parts were then dried overnight and then cured in a Formlabs Form Cure L V1 at 60°C for 1 hour. Parts were sterilized in a sterilization pouch (Plastcareusa, PG-1016) using a vacuum autoclave cycle at 270°F with a 4-minute exposure time, followed by a 30-minute drying process.

### 2.1.2 Environmental control

The environmental control system shown in Figure 1A maintains tissue viability within the bioreactor for long-term experiments (up to 72 hours) while mounted on a confocal microscope. The core of the bioreactor is a 3D printed chamber that isolates the tissue environment. A sealed lid, opened only inside a biosafety cabinet, minimizes exposure to non-sterile conditions and reduces media evaporation. The chamber accommodates a standard 100 mm petri dish which sits on top of a heater (Tokai Hit, TPi-108RH26) which maintains 37°C in the dish. Additionally, the chamber features an inlet for sterile humidified 5% CO which is sourced by an external humidifier (Tokai Hit, TPiDE-HUMID) and a temperature controller with digital gas mixer for CO regulation (Tokai Hit, STXG). For an additional layer of temperature regulation, the full biomechanical bioreactor is enclosed inside a ThermoBox (Tokai Hit, TI2TB-E-BK) which regulates temperature at 37°C around the device. To further minimize media evaporation, the custom chamber includes a reservoir around the petri dish for deionized water (dH O) as a humidity buffer. Our design provides stable temperature control while ensuring a clear optical path for illumination and imaging. In fact, the ThermoPlate has a central opening to provide access for the inverted microscope objective, while the chamber lid has a 35 mm glass coverslip (MatTek, PCS-1.5-35-NON) bonded at the center to allow light transmission to the condenser.

### 2.1.3 Mechanical design

The biomechanical bioreactor was designed to control and measure biomechanical variables (force and displacement) along one axis (uniaxial configuration) or along two axes (biaxial configuration). Computer aided design (CAD) renderings of the device in both configurations are shown in Figure S1 along with an exploded view of the biaxial configuration to highlight individual components. All major components of the device are fixed to a machined aluminum baseplate that fits onto the stage of a Nikon TI2-E inverted microscope. The chamber features two (uniaxial configuration) or four (biaxial configuration) cutouts for mounting the arms that hold the tissue in place. Each arm is mounted onto a linear actuator via a machined aluminum bracket. One arm per axis includes a force transducer positioned between the arm and the linear actuator. Thus, the uniaxial configuration uses two linear actuators and one force transducer, while the biaxial configuration uses four linear actuators and two force transducers (Figure S1).

To impose controlled displacements, linear actuators (Physik Instrumente, M-111.1DG1) were selected to meet the following criteria: to be rated for operation at 37°C, to fit within the microscope enclosure, to be self-locking when unpowered, and to have been previously demonstrated to effectively stretch in vitro cultures within a microscope [20]. To reduce noise in force measurements during stretching, the DC motor variant equipped with a fine threaded spindle was selected. This actuator has a travel range of 15 mm, a resolution of 0.05 µm, and a maximum velocity of 1.5 mm/s. The linear actuators are controlled by a motion controller (Physik Instrumente, C-884.4DC) through a custom Python interface. To measure cell-generated forces, force transducers (Aurora Scientific, 403C) were selected to operate over a narrow force range (±5 mN maximum) while providing high resolution (0.1 µN), thermal stability, and robustness to mechanical overload. To minimize temperature drift, force transducers were pre-warmed in a dedicated incubator at 37°C for 3 days prior to each experiment. The force transducers were connected to the acquisition computer through a two-channel amplifier (Aurora Scientific, 400-AMP-2CH) and a data acquisition (DAQ) board (National Instruments, USB-6003). Force transducer calibration was carried out at 37°C via sequential application of five weights ranging from 40 mg to 350 mg. A linear least squares regression was fit to the force-voltage data generating a slope with units of mg/V that is applied to all experimental data.

### 2.2 Cell culture

NIH/3T3 fibroblasts (ATCC, CRL-1685) were cultured in Dulbecco’s Modified Eagle’s Medium (DMEM, Corning, MT10013CV), supplemented with 10% Fetal Bovine Serum (FBS, ATCC, 30-2020), 1% penicillin-streptomycin (Pen/Strep, ATCC, 30-2300) in humidified incubators at 37°C and 5% CO using T75 flasks (Fisher, 10-126-11). Cultures were monitored regularly by phase contrast microscopy and passaged at 70%-80% confluence. Cells were routinely tested for mycoplasma contamination using the MycoAlert kit (Lonza, LT07-218). Cells between passages 5 and 17 were used for all experiments.

Fibroblast-populated collagen gels were prepared following a modified protocol from Eichinger et al. [18]. To minimize cell proliferation during experiments, cells at 70%-80% confluence were serum-starved for 24 hours by replacing culture media with DMEM supplemented with 0.5% FBS and 1% Pen/Strep. Collagen gels were prepared using previously published protocols [21]. Briefly, acid-solubilized rat tail collagen I (Corning, 354249) was mixed with equal volumes of a neutralizing buffer, consisting of a 100 mM HEPES (Sigma, H3375) solution in 2× PBS adjusted to pH 7.3 via addition of NaOH. Serum-starved fibroblasts were added to achieve a final density of 500,000 cells/mL, and the final collagen concentration was adjusted by adding an experimental media composed of DMEM supplemented with 10% FBS, 10% porcine serum (Gibco, 26250-084), and 1% Pen/Strep. Polydimethylsiloxane (PDMS) molds (described in the supplementary methods) were aligned inside of a 100 mm plastic petri dish and filled with 3 mL (uniaxial configuration) or 6 mL (biaxial configuration) of a 10 mg/mL sterile bovine serum albumin (BSA) solution, which was used to passivate the petri dish for 30 minutes. After washing out the BSA, the dish was positioned within the biomechanical bioreactor and preheated at 37°C for 1 hour. Gel solutions at target concentrations (1.0, 1.5, 2.0, and 3.0 mg/mL) were cast into such pre-warmed PDMS molds and incubated at 37°C for 1 hour. Porous hydrogel inserts, adapted from Eichinger et al. [18] and described in the supplementary methods, were used to anchor the collagen gels to the bioreactor axes during polymerization. Experimental media (25 mL) was then added to the dish to float the gel, and PDMS molds were gently removed with forceps.

### 2.3 Confocal imaging with integrated force monitoring

Once fibroblast-populated collagen gels were polymerized and floated within the biomechanical bioreactor, time-lapse imaging was carried out using a confocal microscope (Nikon, AX-SHS). Confocal reflection microscopy (CRM) and transmitted light detector (TD) images were collected every 30 minutes for 48 hours using a 488 nm laser diode at 1.0% power. A 10× objective with 0.45 numerical aperture was used in combination with a 2.3× zoom and a 2 μs dwell time to image a 512 × 512 pixel area with a resolution of 1.504 μm/pixel. Image stacks spanning from the bottom of the gel to the upper z-limit of the microscope were acquired using ∼2,300 μm z-stacks with a 50 μm step size. Tiling was employed to capture the entire gel width using 16 image tiles with 15% overlap. For the uniaxial configuration, the imaging plane was rotated 45° to capture the compaction perpendicular to the principal axis of the bioreactor. For the biaxial configuration, two tiled z-stacks were captured to measure the horizontal (H, 0°) and vertical (V, 90°) compaction of the central portion of the gel (Figure S8). Force transducer data were acquired through the DAQ and synchronized with image acquisition using the Nikon Elements software. A wiring schematic indicating the approach employed to control linear actuators and record force transducer data during confocal imaging is provided in Figure S6. The analysis pipeline used to assess collagen remodeling and to estimate cross-sectional areas from time-lapse data is described in the supplementary methods.

### 2.4 Immunofluorescence

Fibroblasts were stained for α-smooth muscle actin (α-SMA) to quantify contractile marker expression. After culturing tissue equivalents for a fixed time frame between 0 and 48 hours, the culture media was aspirated from the petri dish, and the tissue equivalent was transferred into a new petri dish containing a PDMS mold with a cutout for a dogbone gel. Samples were fixed with 4% PFA overnight, permeabilized with 0.2% Triton X-100 for 2 hours, and blocked with 2.5% horse serum for 1 hour. Samples were incubated with anti-α-SMA (Biocare, CM001, 1:500), DAPI (Invitrogen, D1306, 1:1000), and Alexa Fluor 568 Phalloidin (Invitrogen, A12380, 1:1000) overnight at 4°C. Subsequently, samples were incubated with Alexa Fluor 647 secondary antibody (Invitrogen, A21422, 1:1000) overnight at 4°C.

Stained samples were imaged using four channels: 405 nm excitation / 420-470 nm emission (DAPI), 561 nm excitation / 570-616 nm emission (Phalloidin), 640 nm excitation / 660-750 nm emission (α-SMA), and TD for DIC imaging. Tiled image stacks of 512 × 512 pixels at a resolution of 3.453 µm/pixel were acquired using a 20× objective with a 10 µm z-step to span the bottom 250 µm of the gel. Cytoplasmic α-SMA intensity in the imaged volume was quantified using a custom ImageJ macro using the following steps. The DAPI and Phalloidin channels were thresholded (intensity range 150-65,535) to generate masks of the nucleus and cytoplasm for each cell. The nuclear mask was subtracted from the cytoplasmic mask to isolate the cytoskeletal region of each cell, which corresponds to the localization of α-SMA expression. This mask was then applied to the α-SMA channel, and the pixel intensity was integrated along the image width to generate an intensity profile for each z-slice. Total α-SMA intensity in the imaged volume was computed by summing the integrated intensities across all z-slices.

### 2.5 Retraction experiments

To validate our stress estimations, a stress-relieving cut was introduced to quantify the elastic recoil due to the active tension generated via fibroblast contraction. After culturing tissue equivalents for 48 hours, gels were cut on one side and retraction was monitored over the course of 60 minutes, with images being captured through a CCD camera mounted on a stereo microscope (Olympus, SZX-ZB7). The percent retraction was calculated as the length of the cut gap divided by the full length of the gel.

### 2.6 Bulk RNA sequencing

Transcriptome-wide gene expression of NIH/3T3 fibroblasts was analyzed by bulk RNA sequencing (RNA-seq) across four collagen concentrations (1.0, 1.5, 2.0, and 3.0 mg/mL), with three biological replicates per group. Following 24 hours of culture in tissue equivalents, cells were isolated via collagenase digestion, RNA was extracted and sequenced, reads were aligned to the mouse reference genome (GRCm39, ENSEMBL v110), and differential expression analysis was performed using PyDESeq2 [22] with the 1.0 mg/mL condition as reference. Pathway overrepresentation analysis was carried out using GSEApy [23] and Enrichr [24] against the “Reactome_Pathways_2024” database [25]. A detailed description of the experimental methods and data analysis pipeline is provided in the supplementary methods.

### 2.7 Statistical analysis

We present experimental data as mean ± standard error of the mean (SEM) or as violin plots with individual data points showing independent measurements. No statistical method was used to predetermine sample size, and investigators were not blinded during experiments or outcome assessment. All statistical analyses were performed using Python 3.13 and SciPy 1.16 [26]. Because of the low sample size, data was assumed to be normal with equal variance. A one-way ANOVA with Tukey HSD post-hoc correction was performed for variables across initial collagen concentrations. Statistically significant values marked: * *p* < 0.05, ** *p* < 0.01, *** *p* < 0.001, **** *p* < 0.0001.

## 3. RESULTS

The novel biomechanical bioreactor design was inspired by the work of Eichinger et al. [18] and built with three key objectives: (1) enabling bioreactor functionality outside a cell culture incubator; (2) miniaturizing the biomechanical testing apparatus; and (3) integrating continuous confocal or multiphoton imaging. Our design comprises a sterile tissue chamber, computer-controlled linear actuators, and high-resolution force transducers mounted on a baseplate that fits within the stage of an inverted confocal microscope (Figure 1). The system operates in two configurations (Figure S1): uniaxial for dogbone tissues and biaxial for cruciform tissues. For high-throughput analyses, we designed simplified bioreactor pods where tissue equivalents can be cultured under uniaxial or biaxial constraints without passive force measurement or active deformation capabilities (Figure S2). Environmental control is maintained through humidified 5% CO and a heating plate in conjunction with a temperature-regulated microscope enclosure (Figure 1A), achieving a stable 37°C set-point within 45 minutes (Figure S3). Porous hydrogel inserts [18] enable efficient transmission of cell-generated forces from collagen gels to force transducers when properly aligned using custom tissue molds (Figures S4 and S5). The platform synchronizes force recording with confocal imaging and supports the application of external deformations via linear actuators (Figure S6), albeit this functionality was not used in this study. Additional technical details are provided in the supplementary methods. In summary, our biomechanical bioreactor integrates sterile tissue culture and confocal imaging with mechanical recording and actuation.

In this work, we used a uniaxial configuration (Figure 1B-C) to culture dogbone-shaped tissue equivalents under static uniaxial constraint (no external stretching) and investigate endogenous stress generation by NIH/3T3 fibroblasts. This cell line was selected for its contractile phenotype [27] and widespread use in validating various tissue engineering platforms [28–35]. Regardless of the specific cell line, it has long been established that in presence of a uniaxial constraint, fibroblasts cultured in collagen collectively develop a tissue-level force (*F*) that integrates individual traction forces (*df*) generated at cell-ECM adhesions (Figure 1D-F). We first measured the force developed by NIH/3T3 fibroblasts cultured over 24 hours (Figure 1G). In line with previous studies [36–39], tissue force generation exhibited three phases: Phase I (lag phase) with negligible force as cells adhere to collagen and extend filopodial processes; Phase II (cell traction) with quasi-linear force increase as cells spread and assume a bipolar morphology; Phase III (cell contraction) where force reaches a steady-state value as cells maintain their morphology and generate nearly constant force under isometric conditions. Each of these steps is accompanied by dramatic remodeling of the collagenous ECM. A key advantage of our platform is that synchronized confocal imaging captures the structural evolution of collagen via the CRM channel while monitoring changes in cell morphology via the TD channel, as tissue equivalents develop endogenous forces (Figure 1H). Starting from an isotropic collagen matrix populated by round cells at 0 hours, the constructs develop into a highly anisotropic collagen network populated by aligned cells at 12 hours, and ultimately into high density tissues when the steady-state force is achieved at 24 hours. Tissue equivalents maintained high viability (Figure S7) and tested negative for mycoplasma, confirming chamber sterility. As proof of concept, we also performed time-lapse acquisitions using biaxially constrained cruciform gels (Figure S8). Overall, our biomechanical bioreactor couples force sensing with real-time microstructural imaging, enabling continuous monitoring of collagen densification and alignment under cell-generated traction forces.

As shown in Movie S1, fibroblasts in uniaxially constrained tissue equivalents align collagen fibers along the direction of tension (X-axis) while compacting the matrix orthogonally (YZ-plane) [40]. Since the true (Cauchy) stress is defined as force applied over a deformed area, the extent and dynamics of this orthogonal matrix compaction dictate stress evolution within tissue equivalents. Therefore, we sought to quantify the temporal evolution of cross-sectional areas in collagen gels from confocal images. However, CRM penetration depth is inherently limited by light scattering and absorption [19], a constraint that worsens as tissue equivalents densify over time. To enable confocal-based area estimation, we used multiphoton microscopy to image the entire cross-sectional area (YZ-plane) from tissue equivalents. A set of uniaxially constrained tissues was imaged at discrete time points using both confocal and multiphoton microscopy (Figure 2). Second harmonic generation (SHG) imaging of collagen revealed that although tissue equivalents had an approximately rectangular cross-section at 0 hours, the bottom surface contracted more than the top surface over time, leading to tissue folding during compaction at 12, 24, and 48 hours (Figure 2A). At the same time, autofluorescence imaging of the metabolic cofactor nicotinamide adenine dinucleotide (NADH) revealed that fibroblasts were evenly distributed at 0 hours but became progressively restricted to a semicircular region at later time points as the gel folded. Indeed, collagen densification and alignment occurred only in cell-populated regions, suggesting that mechanical stress develops exclusively in these areas. Instead, confocal microscopy captured only the bottom and lateral surfaces of the folding gels (Figure 2B). Based on these observations, we developed a semi-automated pipeline that segments the bottom and lateral profiles of the cross section while estimating the top surface using a semicircular arc extrapolation (Figure 2C). This approach enabled frame-by-frame analysis of confocal time-lapse acquisitions to quantify cross-sectional area reduction over time in response to cell-generated forces (Figure S9A). The semicircular arc method reproduced accurately cross-sectional areas measurements conducted using multiphoton microscopy (Figure 2D) and provided improved estimates with respect to manual measurements (Figure S9B). This approach enables, for the first time, continuous quantification of cross-sectional area in tissue equivalents from confocal time-lapse imaging (Figure 2E), a critical capability for estimating the endogenous stress dynamics.

**Figure 2.**
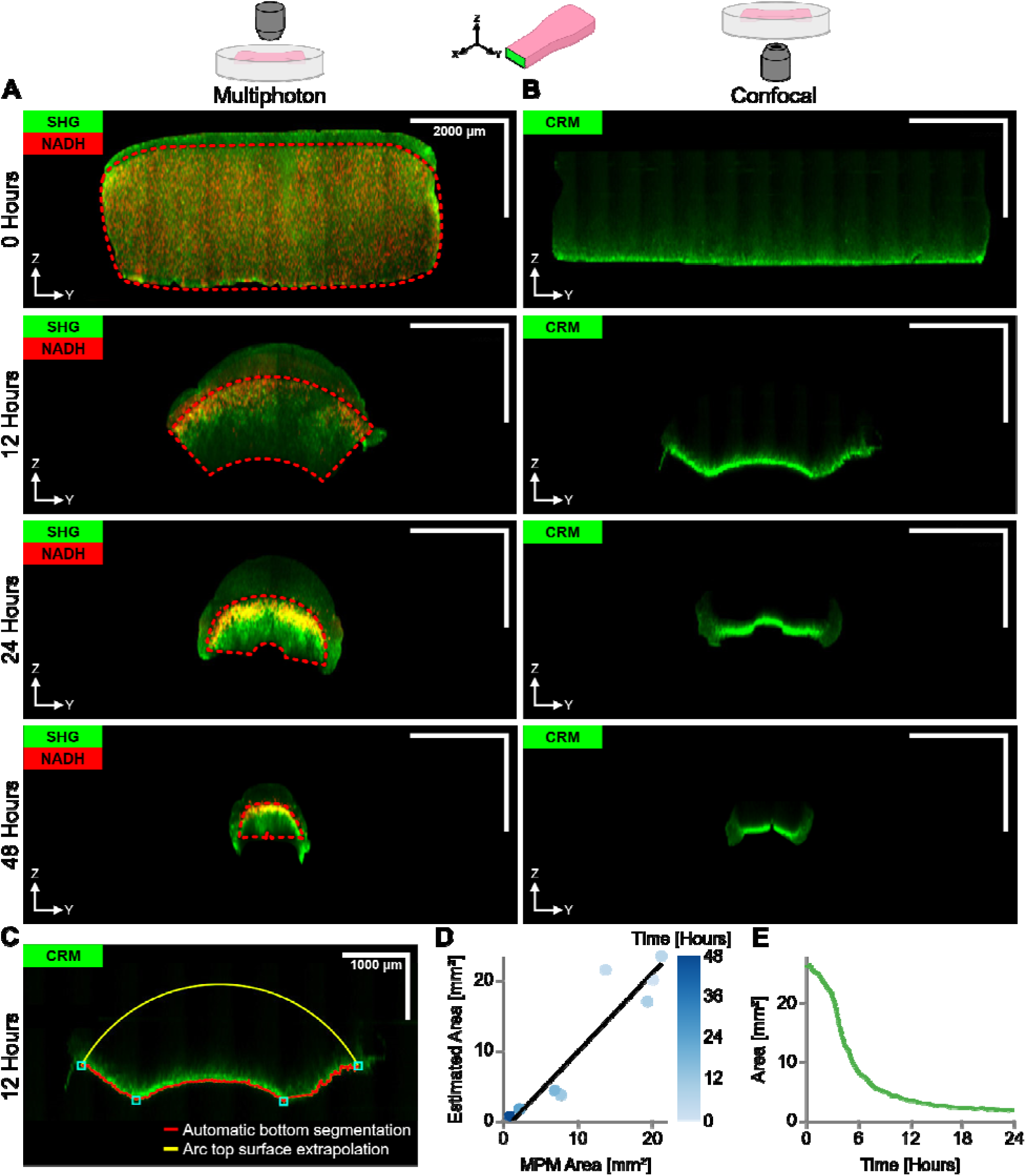
Cross-sectional area quantification and validation in tissue equivalents. (A) Representative multiphoton microscopy (MPM) images of uniaxially constrained gel cross sections (0.8 mg/mL, 500,000 cells/mL) after 0, 12, 24, and 48 hours of remodeling. Collagen was imaged using second harmonic generation (SHG) and fibroblasts were imaged using autofluorescence from nicotinamide adenine dinucleotide (NADH). Scale bars, 2 mm. Cell-generated forces resulted in heterogeneous compaction and folding of the gels over time. Red dashed lines indicate the cell-occupied region within the gel’s cross-section at each time point. (**B**) Paired CRM images of the same samples. The CRM images display strong signal from the bottom of the gel but lack the depth of penetration required to visualize the top surface. (**C**) To quantify the cross-sectional area of the load-bearing fibroblast-populated region from the CRM images, the bottom of the gel was segmented automatically (continuous red line), and the cross-sectional area was estimated using a circular arc to approximate the top surface (continuous yellow line). Scale bars, 1 mm. (**D**) The estimated area from the confocal images using the arc method tracks linearly with the manually segmented MPM cross-sectional area. R² = 0.869. (**E**) Estimated cross-sectional area over time corresponding to the force recording shown in Figure 1G, showing the dynamics of tissue compaction under the action of cell-generated forces.

Following prior work on free-floating [15] and anchored [41] gels, we used our system to test the hypothesis that fibroblasts achieve tensional homeostasis by maintaining a nearly constant state of stress. We cultured NIH/3T3 fibroblasts at a fixed cell density of 500,000 cells/mL in gels with increasing initial collagen concentrations (*p*_0_ = 1.0, 1.5, 2.0, 3.0 mg/mL). This range encompasses low to high collagen concentrations commonly employed in tissue equivalent research. After 48 hours, macroscopic gel compaction was greater at lower *p*_0_ (Figure 3A). Cell-generated forces increased over time, with different initial collagen concentrations exhibiting distinct kinetics. For the 1.0-2.0 mg/mL range, force measurements reached a steady-state value between 24 and 48 hours, with final values increasing with *p*_0_ (Figure 3B), consistent with previous reports [16,18]. Conversely, force measurements in 3.0 mg/mL gels showed delayed kinetics and did not reach steady-state by 48 hours. Estimated cross-sectional areas were initially indistinguishable across groups and decreased nearly exponentially over time (Figure 3C). The extent by which the cross-sectional area decreased was found to vary as a function of the initial collagen concentration: 1.0 mg/mL gels underwent greater compaction, displaying lower cross-sectional areas at 24 and 48 hours, while 3.0 mg/mL gels underwent less compaction, maintaining higher cross-sectional areas at both time points (Movie S2). By independently measuring the tissue force (*F*) and cross-sectional area (*a*), our platform enables estimation of the average Cauchy stress (*a*= *F/a*) over time (Figure 3D). In contrast to force and cross-sectional area, which plateaued in most groups, the Cauchy stress increased monotonically over time without reaching a steady-state value within 48 hours. More importantly, stress was not preserved across groups but decreased with increasing *p*_0_: at 48 hours, the Cauchy stress was highest in 1.0 mg/mL gels compared to all other concentrations (Figure 3D). To rule out potential errors from our force and our cross-sectional area measurements, we independently released the active stress developed by fibroblasts upon contraction. After culturing uniaxially constrained tissue equivalents in bioreactor pods for 48 hours, we introduced a stress-relieving cut (Figure 3E) and quantified the retraction dynamics (Figure 3F). Gels retracted rapidly upon cutting, consistent with a nearly instantaneous release of active stress (Movie S3). Critically, the extent of retraction mirrored the trends observed in the Cauchy stress: 1.0 mg/mL gels displayed the highest retraction (∼25%) while 3.0 mg/mL gels showed minimal retraction (∼1.5%). Taken together, these results challenge the assumption that mechanical stress is the homeostatic target in fibroblast-populated tissue constructs.

**Figure 3.**
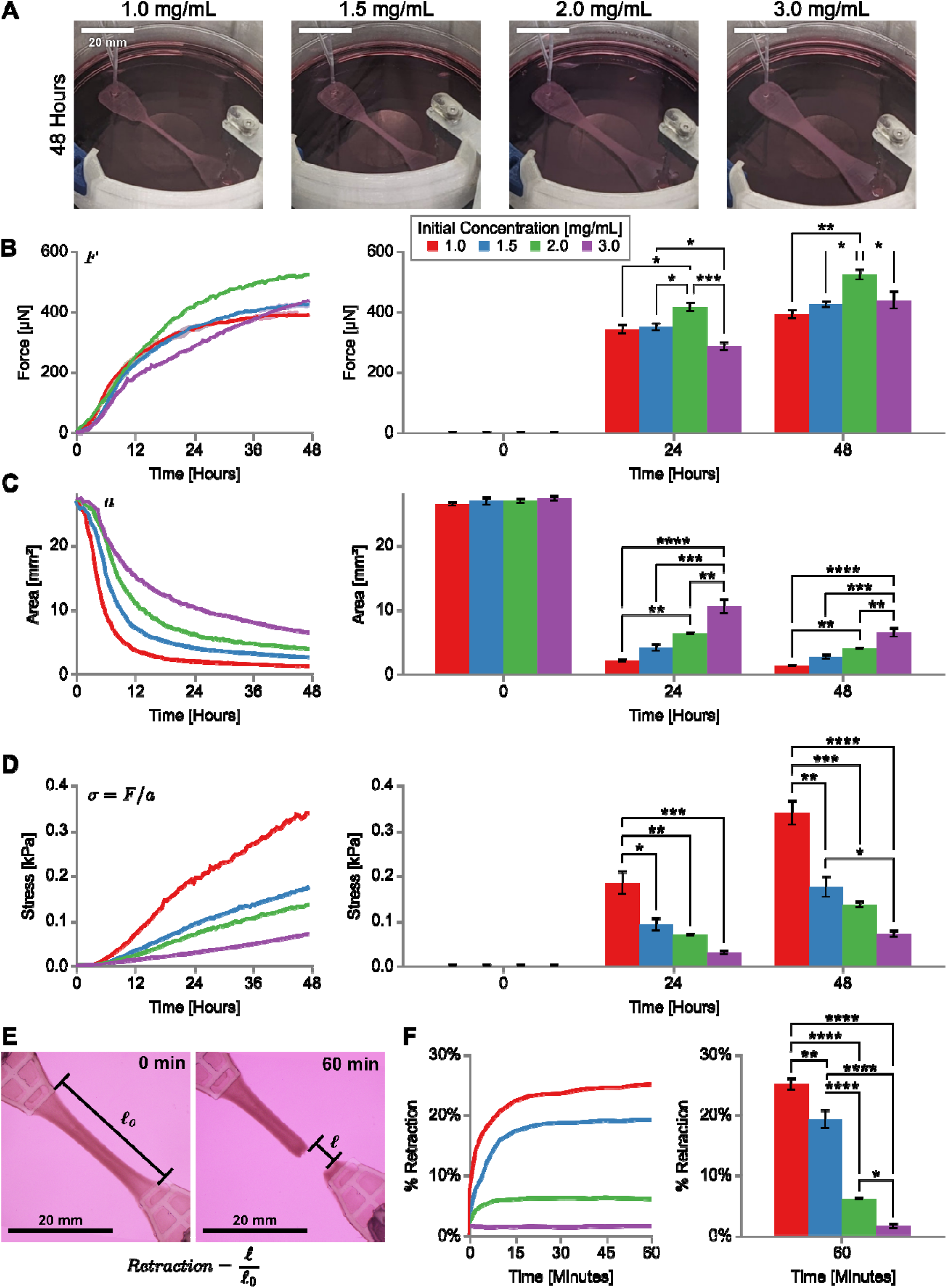
Mechanical stress is not maintained across collagen concentrations. (**A**) NIH/3T3 fibroblasts at a fixed cell density of 500,000 cells/mL were cultured in gels of varying collagen concentrations: 1.0 mg/mL (n=4), 1.5 mg/mL (n=3), 2.0 mg/mL (n=3), and 3.0 mg/mL (n=3). (A) Macroscopic images of uniaxial gel compaction after 48 hours of remodeling for each concentration reveal greater compaction in lower initial concentration gels. Scale bar, 20 mm. (B) Cell-generated forces reach a plateau after 48 hours, where gels with higher initial collagen concentration (in the range 1.0 – 2.0 mg/mL) reach higher steady-state forces, while the 3.0 mg/mL gels lag behind. (**C**) The cross-sectional area decreases with time in a concentration-dependent manner, where gels with higher initial concentrations display a larger cross-sectional area consistent with a lower degree of compaction. (**D**) Contrary to our initial hypothesis, mechanical stress (calculated dividing the current force by the current cross-sectional area) is not maintained constant across collagen concentrations but rather is greater for lower initial collagen concentrations. It should be noted that stress does not reach a steady-state value by 48 hours. (**E**) As an independent validation for the observed trends in stress, the gel retraction *(ℓ⁄ℓ*_0_) after 48 hours of remodeling was measured. Shown is a representative image of a 1.5 mg/mL gel before and after introducing a cut perpendicular to the direction of force generation. (**F**) After such stress-relieving cut, gel retraction was monitored and quantified over 60 minutes for the four concentrations, 1.0 mg/mL (n=5), 1.5 mg/mL (n=3), 2.0 mg/mL (n=3), and 3.0 mg/mL (n=4). It was found that gels with lower initial concentrations exhibit greater percent retraction, mirroring the trends in mechanical stress observed from the bioreactor experiment. These observations support the idea that mechanical stress does not represent the homeostatic target over the short-term (48 hours). Data in all time courses and bar plots are shown as mean ± SEM. Statistical significance was assessed using a one-way ANOVA with equal variance assumption and a Tukey post-hoc test; * *p<0.05*, ** *p<0.01*, *** *p<0.001*, **** *p<0.0001*.

The observation that mechanical stress does not reach a steady-state value by 48 hours while varying dramatically across groups prompted us to explore other physical quantities. We used a combination of optical clearing, multiphoton microscopy, and 3D image analysis to quantify cell nuclei distribution and collagen volume in the central regions of tissue equivalents (Figure 4A). Although all groups had the same initial cell density, tissues with *p*_0_ *=*1.0 mg/mL reached a final cell density that was twice that of the other groups at 48 hours (Figure 4B). This finding indicated that tissue equivalents with initially low collagen concentration display elevated cell density due to increased tissue compaction. The true collagen density in the remodeled tissue equivalents can be computed as *p* = *p*_0_ *· (a*_0_*/a)*, where *a*_0_ indicates the initial cross-sectional area (identical across groups) and *a* represents the current cross-sectional area (estimated from confocal imaging). We found that the true collagen density *p* at 24 and 48 hours decreased dramatically with increasing initial collagen concentration *p*_0_ (Figure 4C). This means that gels with initial collagen density of *p*_0_ *=* 1.0 mg/mL underwent the highest collagen densification, reaching a true collagen density of *p* ≅ 25 mg/mL after 48 hours, a 25-fold increase driven by tissue compaction. On the other hand, gels with the highest initial collagen density of *p*_0_ *=* 3.0 mg/mL underwent the lowest collagen densification, reaching a true collagen density of *p* ≅ 13 mg/mL after 48 hours, slightly more than a 4-fold increase. We found that the Cauchy stress scaled inversely with the initial collagen concentration (*σ ∞ 1⁄p*_0_) at all time points (Figure S10A). At the same time, the Cauchy stress follows a power-law relationship with the true collagen density (*σ ∞ p*^0^) and the quality of the power-law fit improves progressively over time (Figure S10B). Based on these observations, we explored whether active stress generation depends on the true collagen density by plotting *σ(t)* against *p(t)* for each experimental group (Figure 4D). We found that the active stress generation displays a strong dependency on the true collagen density, and we interpreted this relationship using the following power-law description: *σ* = *ap*^b^, where each initial collagen concentration *p* may display different parameters. The proposed power-law fits the experimental data reasonably well (Figure 4D), with the parameters *a* and *b* displaying consistent trends with respect to the initial collagen concentration *p*_0_. Specifically, we found that *a* decreases while *b* increases with *p*_0_ (Figure 4E-F). We interpreted these trends as follows: low density tissue equivalents (1 mg/mL) are very sensitive and generate active stress in an almost linear fashion in response to collagen densification. Instead, high density tissue equivalents (3 mg/mL) are less sensitive and generate active stress in an almost quadratic fashion in response to collagen densification. Therefore, active stress generation by fibroblasts depends directly on the true collagen density, which is in turn regulated by local cell contractility.

**Figure 4.**
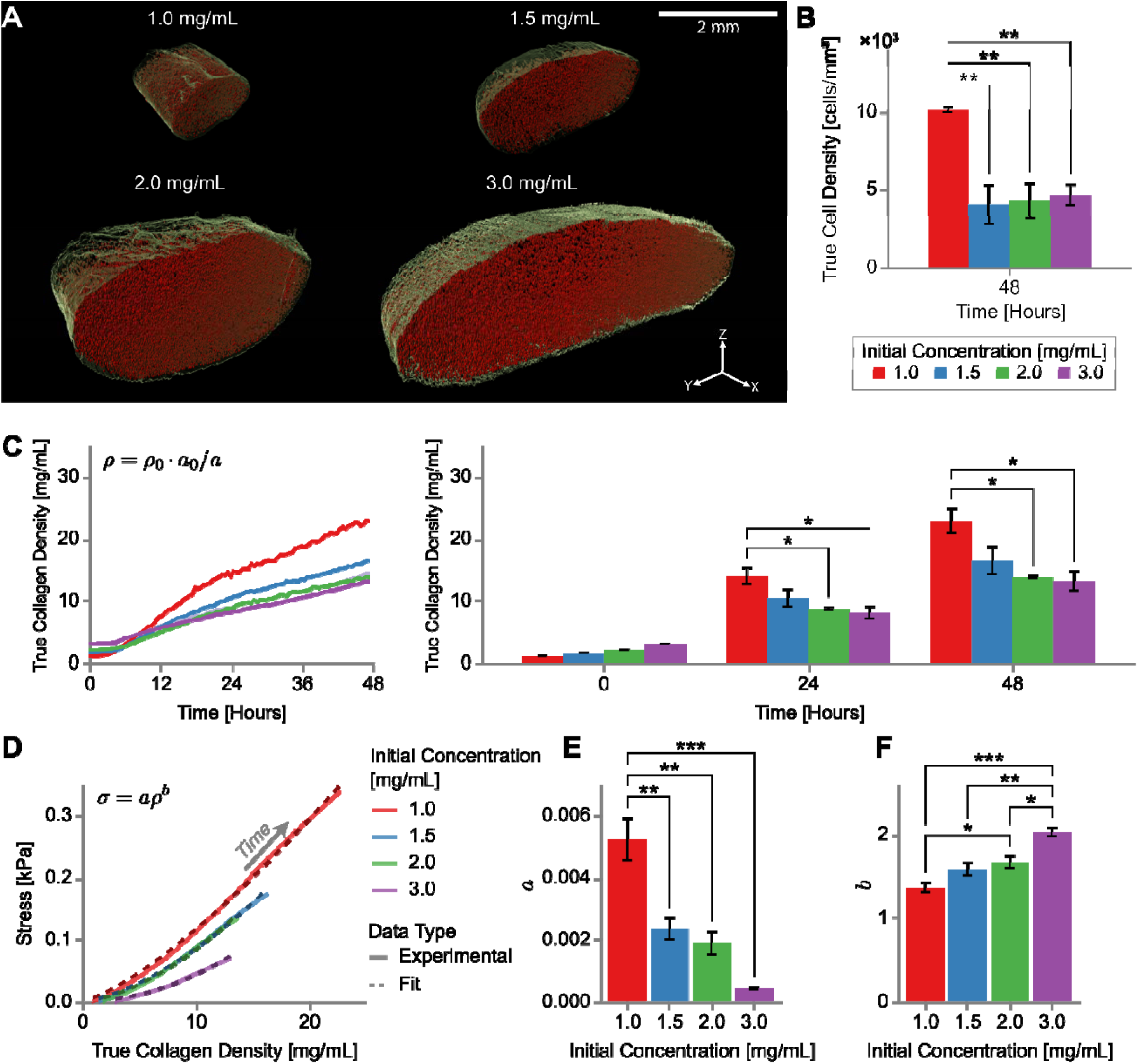
Collagen densification is associated with active stress generation.**3D renderings of** the central region from representative tissue equivalents. Multiphoton images were analyzed using an Imaris workflow that isolates the DAPI-stained cell nuclei (red dots) via spot detection and the SHG-positive collagen volume (green surface) via surface segmentation. (**B**) The density of cells after 48 hours of remodeling was calculated by dividing the number of detected nuclei by the remodeled collagen volume (n=3 for all concentrations). (**C**) The true collagen density was calculated by scaling the initial collagen concentration (*p*_0_) by the ratio of current (*a*) and initial (*a*_0_) cross-sectional areas. This calculation reveals that collagen gels with the lowest initial collagen concentration (*p*_0_ = 1.0 mg/mL) display the highest densification both at 24 and 48 hours. (**D**) Active stress generation *a* displays a power-law dependency on the true collagen density *p* as shown by average *a- p* data plotted for each group (continuous line) and fitted using the equation *a= ap*^b^ (dashed line). The power-law parameters (**E**) *a* and (**F**) *b* display opposite trends as a function of the initial collagen concentration, consistent with the Cauchy stress displaying increased sensitivity and reduced nonlinearity for *p*_0_ = 1.0 mg/mL. Data in all time courses and bar plots are shown as mean ± SEM. Statistical significance was assessed using a one-way ANOVA with equal variance assumption and a Tukey post-hoc test; * *p<0.05*, ** *p<0.01*, *** *p<0.001*, **** *p<0.0001*.

To validate the predicted trends in true collagen density *p*, we analyzed the temporal evolution of collagen remodeling using dynamic CRM imaging data (Movie S4). Fibroblasts remodeled collagen in a concentration-dependent manner despite identical initial cell density and morphology (Figure 5A). Polar histograms from a segmentation analysis conducted using the Ridge Detection algorithm [42] revealed that collagen fibers, initially randomly oriented, progressively realigned along the direction of force generation at rates that depended on the initial collagen concentration (Figure S11A). Indeed, fibers in 1.0 mg/mL gels aligned earlier and more extensively than those in 3.0 mg/mL gels. Fitting the percentage of aligned fibers with a logistic function (Figure S11B) allowed us to quantify the alignment time constant for each initial collagen concentration (Figure 5B). A smaller time constant indicates earlier alignment (for 1.0 mg/mL gels) while a higher time constant suggests a later alignment (for 3.0 mg/mL gels). Additionally, we exploited the fact that the CRM intensity is proportional to the local collagen density [43,44]. We found that CRM intensity quantified in the central regions of tissue equivalents followed the expected collagen density trends: initially lower for 1.0 mg/mL gels, this signal increased dramatically over time, ultimately exceeding all other groups (Figure S11C). The inverse relationship between CRM intensity at 24 hours and the initial collagen concentration *p*_0_ (Figure 5C) corroborates our collagen density calculations (Figure 4C). Collectively, analysis of collagen remodeling dynamics revealed that, despite generating the lowest forces, tissue equivalents with the lowest initial collagen concentration (1.0 mg/mL) develop the highest stresses due to higher densification and earlier alignment of collagen fibers.

**Figure 5.**
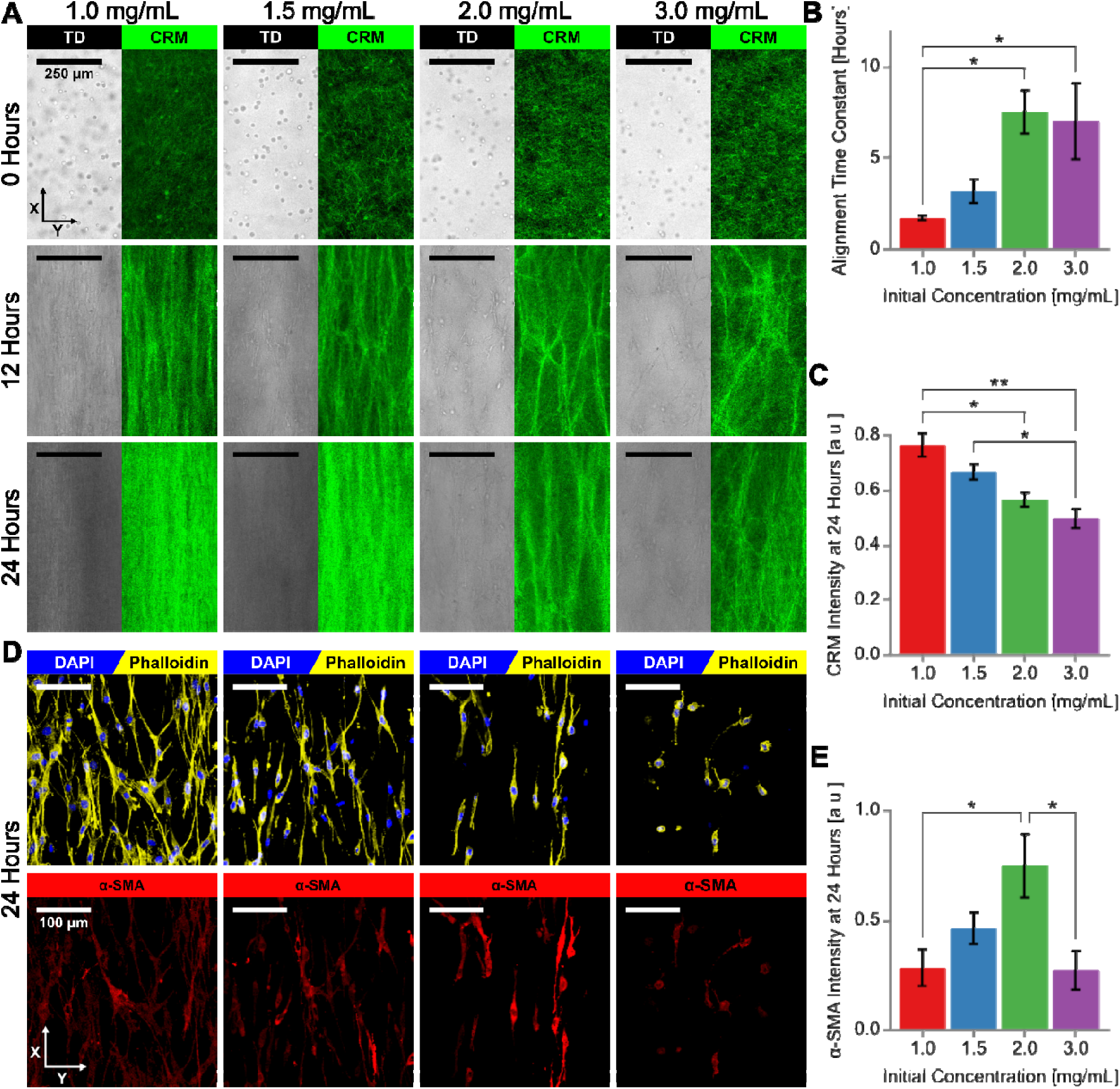
Local assessment of collagen densification and cell contractility. (A) Representative DIC and CRM images for the four initial collagen concentrations at 0, 12, and 24 hours display earlier alignment and increased densification of collagen fibers in lower initial concentration gels. (**B**) Quantification of the alignment time constant () from a sigmoid fit for the percentage of aligned fibers (i.e., fibers aligned along the principal axis of the bioreactor) reveals that gels with higher initial concentrations display higher values consistent with a delayed alignment. (**C**) Average CRM signal intensity in the cell-populated region of gel quantified at 24 hours parallels the trend in true collagen density ρ from Figure 4C. Group sizes for CRM quantifications: 1.0 mg/mL (n=4), 1.5 mg/mL (n=3), 2.0 mg/mL (n=3), and 3.0 mg/mL (n=3). (**D**) Representative immunofluorescence images showing staining with DAPI and phalloidin (top row) as well as α-SMA (bottom row) for the four initial collagen concentrations at 24 hours display a biphasic expression of the contractile marker. It should be noted how fibroblasts display morphological changes as a function of the initial collagen concentration, with bipolar morphologies in the range 1.0 – 2.0 mg/mL and a stellate morphology in 3.0 mg/mL. (**E**) Quantification of total α-SMA intensity from the cell cytoplasm at 24 hours parallels the trend in cell contractile force from Figure 3B. Group sizes for α-SMA quantifications: 1.0 mg/mL (n=3), 1.5 mg/mL (n=5), 2.0 mg/mL (n=4), and 3.0 mg/mL (n=4). Data in all bar plots are shown as mean ± SEM. Statistical significance was assessed using a one-way ANOVA with equal variance assumption and a Tukey post-hoc test; * *p<0.05*, ** *p<0.01*, *** *p<0.001*, **** *p<0.0001*.

To validate the observed trends in cell contractile force *F*, we quantified α-smooth muscle actin (α-SMA) expression using immunofluorescence. We developed a pipeline to measure total cytoplasmic α-SMA intensity in tissue equivalents (Figure S12A-B) and used it to track α-SMA expression over time in 1.0 mg/mL gels (Figure S12C). Total α-SMA intensity was initially low and increased sigmoidally over time, reaching a plateau around 12 hours, notably preceding the force increase measured in bioreactor experiments (Figure S12D). This temporal sequence suggests a causal relationship between α-SMA expression and force generation. Next, we quantified total α-SMA intensity in gels with graded collagen concentrations fixed at 24 hours (Figure 5D). Consistent with the observed pattern in contractile force (Figure 3B), α-SMA intensity increased across the 1.0-2.0 mg/mL range but decreased at 3.0 mg/mL (Figure 5E). Importantly, a lower α-SMA intensity was accompanied by a morphological switch from bipolar to stellate morphology at 3.0 mg/mL. Overall, these data confirm the biphasic trend in cellular contractility observed across the range of initial collagen concentrations considered herein.

The observed trends indicate that tensional homeostasis may emerge from a balance between cell contractility and collagen densification: fibroblasts generate higher contractile forces when high density matrices resist compaction and lower contractile forces when low density matrices compact more readily. This homeostatic balance holds across the 1.0-2.0 mg/mL range but breaks down at 3.0 mg/mL, where fibroblasts display reduced α-SMA expression, diminished contractile force, and limited collagen densification. To investigate the molecular basis of this disrupted phenotype, we performed bulk RNA sequencing on cells isolated after 24 hours in culture (Figure 6). Principal Component Analysis (PCA) revealed modest transcriptional differences between fibroblasts cultured in 1.0, 1.5, and 2.0 mg/mL, but a more distinct transcriptional profile for fibroblasts cultured in 3.0 mg/mL (Figure 6A).

**Figure 6.**
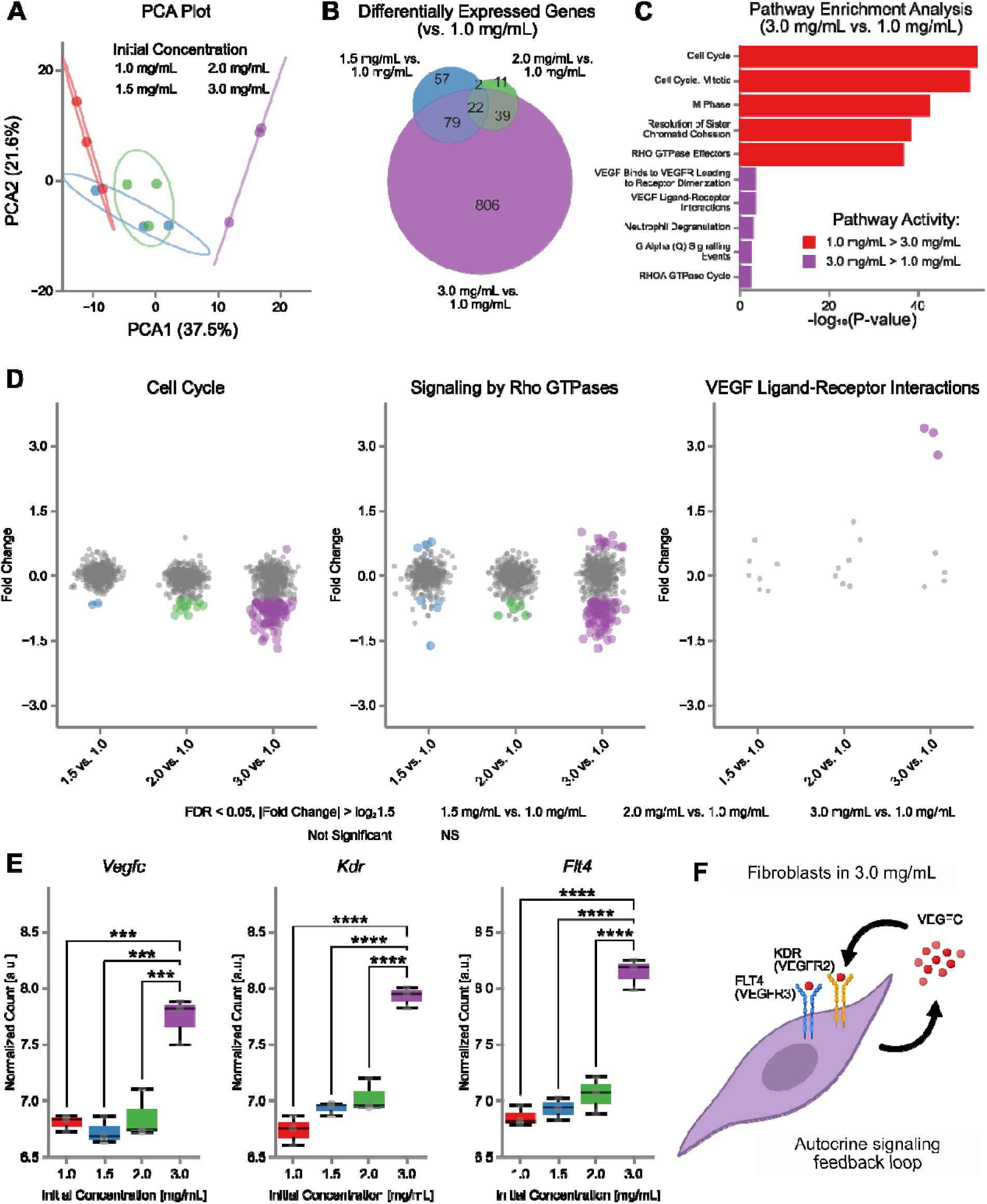
Transcriptional changes associated with disrupted tensional homeostasis. (A) Principal component analysis (PCA) of variance-stabilized bulk RNA sequencing data from fibroblasts cultured for 24 hours in graded collagen concentrations: 1.0 mg/mL (n=3), 1.5 mg/mL (n=3), 2.0 mg/mL (n=3), and 3.0 mg/mL (n=3). (**B**) Differentially expressed genes (DEGs) [fold change (FC) > 1.5 and false discovery rate (FDR)-adjusted P < 0.05] compared to 1.0 mg/mL. (**C**) Top enriched Reactome [25] pathways (FDR < 0.05) for significantly up- and downregulated genes in 3.0 mg/mL vs. 1.0 mg/mL gels. (**D**) Gene expression in overrepresented Reactome pathways. Data are presented as violin plots with data points indicating individual genes. Colored dots indicate DEGs in each pathway that are significant (FDR < 0.05 and FC > 1.5) for a specific comparison, while gray dots indicate genes that are not significant (NS). Violin distributions reflect only statistically significant genes. (**E**) Box plots of differential overexpressed genes in the “VEGF Ligand-Receptor Interactions” pathway for 3.0 mg/mL vs. 1.0 mg/mL. Genes encoding for both growth factor (*Vegfc*) and its receptors (*Kdr* and *Flt4*) are upregulated in fibroblasts cultured in 3.0 mg/mL collagen. (**F**) Model depicting a fibroblast in 3.0 mg/mL collagen establishing an autocrine signaling loop to promote cell survival. Statistical significance was assessed using a one-way ANOVA with equal variance assumption and a Tukey post-hoc test; * *p<0.05*, ** *p<0.01*, *** *p<0.001*, **** *p<0.0001*.

Differential expression analysis identified 946 differentially expressed genes (DEGs) in the 3.0 mg/mL vs. 1.0 mg/mL comparison, compared to 160 DEGs for 1.5 mg/mL vs. 1.0 mg/mL, and 74 DEGs for 2.0 mg/mL vs. 1.0 mg/mL (Figure 6B). Volcano plots for all comparisons confirmed that the largest number of DEGs was specific to the 3.0 mg/mL vs. 1.0 mg/mL comparison (Figure S13). We therefore conducted pathway overexpression analysis using the Reactome database to identify differentially regulated pathways in the 3.0 mg/mL vs. 1.0 mg/mL comparison (Figure 6C). Among the top enriched pathways, downregulated pathways (more active in 1.0 mg/mL) included cell cycle and signaling by Rho GTPases. In contrast, upregulated pathways (more active at 3.0 mg/mL) notably involved VEGF signaling (Figure 6C). To visualize these trends across all comparisons, we plotted the fold change (FC) for genes in three selected Reactome pathways: cell cycle, signaling by Rho GTPases, and VEGF ligand-receptor interactions (Figure 6D). Heatmaps of significant DEGs in these pathways revealed a gradual transition in transcriptional profiles with increasing collagen concentration (Figure S14). Both cell cycle and signaling by Rho GTPases displayed progressive downregulation across the range of initial collagen concentrations (1.5, 2.0, 3.0 mg/mL compared to 1.0 mg/mL). However, the VEGF ligand-receptor interactions pathway showed no differentially expressed genes reaching statistical significance for the 1.5 mg/mL vs. 1.0 mg/mL and 2.0 mg/mL vs. 1.0 mg/mL comparisons. Instead, this pathway displayed three significantly upregulated genes for the 3.0 mg/mL vs. 1.0 mg/mL comparison. Figure 6E shows the expression of these three upregulated genes *Vegfc*, *Kdr*, and *Flt4*. Strikingly, *Vegfc* encodes vascular endothelial growth factor C (VEGFC), while *Kdr* and *Flt4* encode vascular endothelial growth factor receptor 2 (VEGFR2) and vascular endothelial growth factor receptor 3 (VEGFR3), respectively. The coordinated upregulation of both the growth factor (VEGFC) and its receptors (VEGFR2 and VEGFR3) suggests that fibroblasts at 3.0 mg/mL can establish an autocrine signaling loop to promote cell survival (Figure 6F). We examined the expression of key genes across the four initial collagen concentrations (Figure S15) and found that fibroblasts at 3.0 mg/mL upregulate the expression of planar cell polarity genes (*Celsr1* and *Dchs1*) and vascular signaling molecules (*Gdf2*, *Sox18*, *Tek*, *Eng*). These results indicate that fibroblasts maintain tensional homeostasis (1.0-2.0 mg/mL) largely through post-translational mechanisms that involve minimal changes in gene expression, but resort to transcriptional reprogramming in dense matrix microenvironments (3.0 mg/mL), leading to autocrine signaling through VEGFC that promotes cell survival.

## 4. DISCUSSION

The ability of tissues to maintain their mechanical properties within a physiologically optimal range is referred to as mechanical homeostasis, while maintenance of such a homeostatic state by cells through the generation of tensional forces is called tensional homeostasis [8]. Stromal fibroblasts are the primary cell type involved in the regulation of tensional homeostasis through various ECM remodeling mechanisms, including ECM deposition, degradation, cross-linking, as well as force-mediated densification and alignment [45]. The latter mechanism is particularly relevant during the activation of stromal fibroblasts into contractile myofibroblasts, which leads to an increase in mechanical tension within the connective tissue compartment [46].

In turn, mechanical tension is known to drive early and persistent myofibroblast differentiation both in vivo [47] and in vitro [48]. Therefore, mechanical tension is both a cause and a consequence of myofibroblast activation, which suggests that load-bearing tissues adapt their mechanical properties via a homeostatic feedback loop involving fibroblast-driven ECM remodeling [49,50]. The exact mechanisms regulating this homeostatic control system are largely unknown, even though they appear to regulate both normal tissue function [51] and the progression of pathological conditions, such as fibrosis and cancer [52]. In this work, we presented a novel platform – a biomechanical bioreactor that integrates dynamic force sensing with continuous microstructural imaging – and used it to examine the feedback control system regulating tensional homeostasis. We identified a balance between cell contractility and collagen densification as a potential homeostatic set-point for fibroblasts based on the underlying physics and biology (Figure 7).

**Figure 7.**
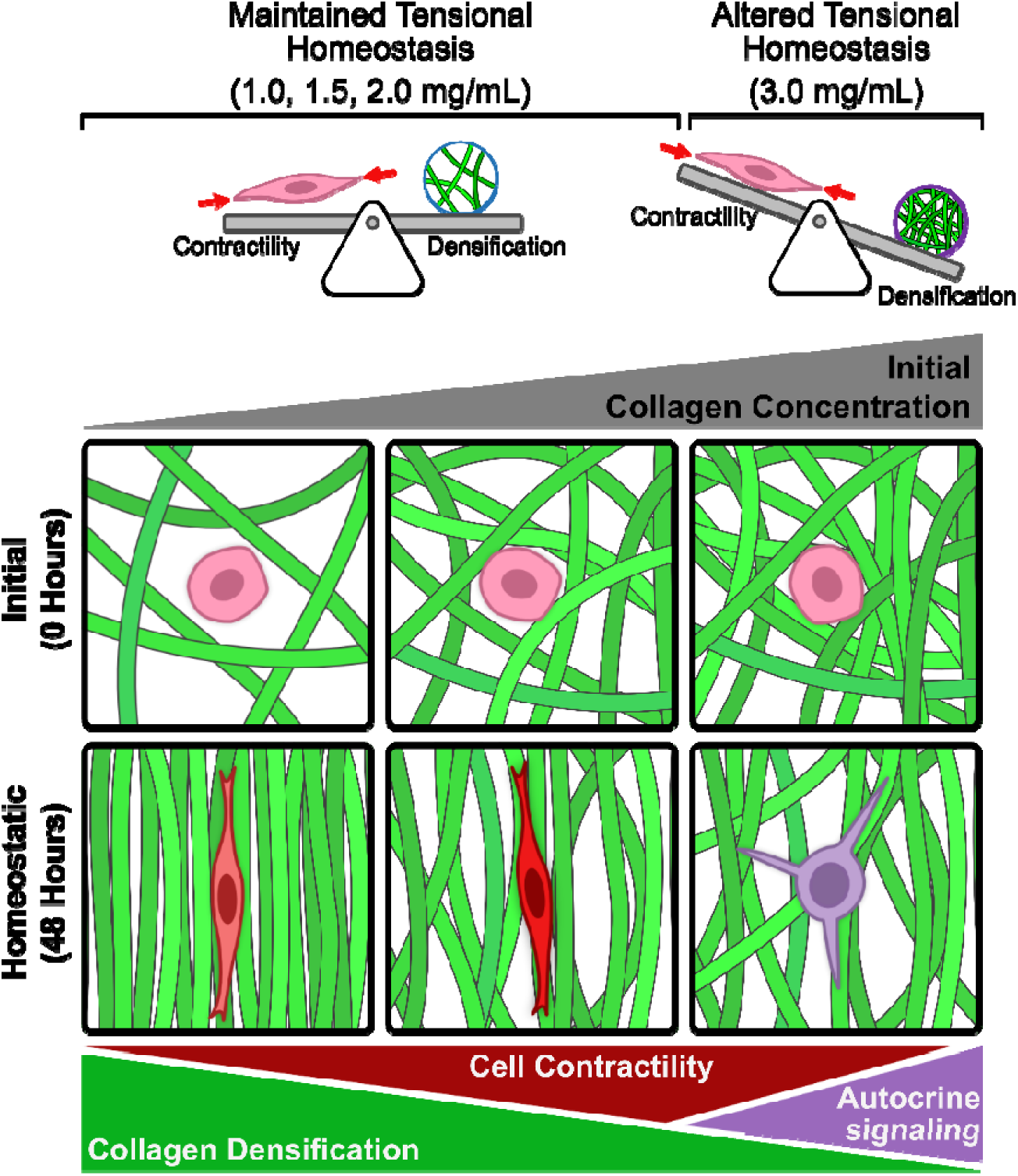
Tensional homeostasis emerges from a balance between cell contractility and collagen densification. Using our novel biomechanical bioreactor, we found that mechanical stress is not maintained across different collagen concentrations over the course of 48 hours. Instead, a balance between cell contractility and collagen densification is maintained nearly constant for initial collagen concentrations between 1.0 mg/mL and 2.0 mg/mL. This homeostatic balance is disrupted in 3.0 mg/mL collagen, where fibroblasts exhibit reduced collagen densification, decreased α-SMA expression, and a transcriptional profile consistent with an autocrine signaling loop.

To identify the homeostatic set-point regulating the complex cell-ECM interactions underlying tensional homeostasis, one needs to monitor dynamically the macroscopic tissue-level force and geometry as well as the microscopic organization of cells and the ECM. Over the years, several engineering platforms have been developed and have been used to explore tensional homeostasis at multiple length scales. For an overview of the existing platforms, the interested reader is referred to the supplementary discussion. However, since existing platforms are limited in their ability to collect both force and imaging data dynamically [16–19,30], here we proposed an integrated bioreactor design that combines simultaneous biomechanical, microstructural, and geometrical measurements. Our goal was to culture macroscopic tissue equivalents under uniaxial or biaxial loads, while enabling simultaneous force and structural measurements. The resulting bioreactor enables simultaneous monitoring of collagen microstructure and tissue-level forces (Figure 1). The magnitude of forces generated by NIH/3T3 fibroblasts in our system is in agreement with those previously reported by Eichinger et al. [18], although in their system steady-state forces were achieved earlier (12-24 hours) than in our system (24-48 hours). Confocal imaging allowed us to make the first dynamic measurements of cross-sectional area evolution in collagen gels during fibroblast-mediated remodeling (Figure 2). Even though our platform can be used in uniaxial or biaxial configurations, we found that uniaxial contraction leads to tissue folding, which reveals heterogeneous tensions within the cross-sectional area (Movies S1-S2). Even though fibroblasts appear initially distributed homogeneously throughout the rectangular cross-section, over the course of 48 hours, the bottom surface contracts heavily, pulling down the sides and forming an arc shape (Figure 2A). This behavior may simply be caused by a higher cell density at the bottom of the gel due to cell sedimentation. However, our analysis of α-SMA expression suggests that contractile fibroblasts concentrate at the gel periphery (Figure S12), including the bottom and side surfaces (the top surface was inaccessible by confocal microscopy). This suggests that, similar to observations in free-floating tissue equivalents [11], contractile fibroblasts may segregate toward the outer surface of uniaxially constrained tissue equivalents where they can effectively generate tensile forces. The spatial distribution of contractile fibroblasts in tissue equivalents and its relation to tensional homeostasis deserve further investigation.

Tissue equivalents are complex materials. The dramatic reduction in cross-sectional area reveals highly compressible behavior, consistent with the large reduction in thickness observed when collagen gels are exposed to external loads [19,53,54]. Moreover, the material properties of tissue equivalents evolve over time due to cell-generated contraction and collagen remodeling. Contractility represents an active contribution to the mechanics of tissue equivalents, following a Hill-type material behavior and thereby increasing the measured tissue stiffness [55]. In contrast, collagen remodeling represents a passive contribution that bears an increasingly higher fraction of the endogenously generated tension over time [39]. Furthermore, collagen alignment is associated with anisotropic stiffening of tissue equivalents along the direction of the imposed constraint [56]. Anisotropic stresses have been recently shown to sustain the phenotypic transition of fibroblasts into myofibroblasts [35]. Although some of these behaviors are well characterized, stress estimations within tissue equivalents are not common and thus far have been conducted only during mechanical testing [19]. Our bioreactor platform enables dynamic assessments of the total (active and passive) Cauchy stress *a* through direct measurements of the contractile force *F* and of the remodeled cross-sectional area *a* (Figure 3A-C). We found that the total stress developed by tissue equivalents is not maintained constant across different collagen microenvironments but rather depends on the initial collagen concentration *p*_0_: while all tissue equivalents are initially stress-free, over time the Cauchy stress reaches higher values for lower *p*_0_ and lower values for higher *p*_0_ (Figure 3D). We confirmed this crucial finding by measuring the extent of retraction in response to a stress-relieving cut (Figure 3E-F). While our measurements only capture the average Cauchy stress from macroscopic measurements, it is worth noting that our results mirror the findings obtained using microfabricated systems [30], where mechanical stress also decreases with increasing collagen density. Dynamic microstructural quantifications of collagen density and alignment (Figure 5A-C) allowed us to interpret the increased stress in tissues with lower *p*_0_ as resulting from higher densification and earlier alignment of collagen fibers (Figures 5 and S10).

The concept of homeostatic stress is rooted in early experimental observations [57] and is central to our understanding of tissue growth and remodeling [58]. In theoretical models, tissue growth and remodeling is triggered by a departure from the homeostatic stress [59]. While the existence of homeostatic stress continues to be revealed at the tissue-level [60], our work suggests that the cell-mediated mechanisms by which fibroblasts establish a favorable microenvironment in connective tissues are not aimed at maintaining a constant level of stress. Since neither force nor stress are preserved across different initial collagen concentrations, it is quite compelling to ask which metric, if any, may regulate the development of tensional homeostasis. Prior experimental and modeling work has indeed suggested that cells aim to restore the contractile forces exerted on the surrounding ECM [61]. However, the study by Eichinger et al. [61] considered mechanical perturbations at a fixed collagen concentration as opposed to our analysis, in which we examined a range of collagen concentrations without applying mechanical perturbations. Our data suggest that fibroblasts may attempt to preserve a balance between cell contractility and matrix densification. If true, this finding implies that tensional homeostasis arises from a balance between cell contractility (biological factor) and collagen densification (physical factor). We demonstrated the relevance of the proposed homeostatic balance by showing that the contractile force exerted by cells follows the trend observed in α-SMA expression [62], while the true collagen density *p* follows the trend observed in CRM intensity (Figure 5). The increase in contractility and the parallel decrease in collagen densification observed between 1.0 mg/mL and 2.0 mg/mL involved modest transcriptional changes (Figure 6), suggesting that tensional homeostasis can be modulated by cells at the protein level without requiring changes in gene expression. Hence, while the true nature of the homeostatic set-point remains to be identified, our work indicates that tensional homeostasis may develop through a balance between cell contractility and collagen densification (Figure 7). The mechanistic identification of competing homeostatic metrics warrants further investigation and will likely require a combination of theoretical analyses and experimental validations.

A notable exception to the homeostatic scenario presented in Figure 7 is provided by tissue equivalents cultured in high initial collagen concentrations (*p*_0_ *=* 3.0 mg/mL). To our knowledge, this represents the first study examining tensional homeostasis in collagen microenvironments at such elevated densities. The study by Pizzo et al. [28] used anchored tissue equivalents to show that fibroblasts cultured in 3.0 mg/mL preserve a rounder morphology with an increased number of filopodial projections while displaying reduced ECM remodeling and lower proliferation compared to fibroblasts cultured in 1.0 mg/mL. Consistent with this work, we also found that by 24 hours fibroblasts in 3.0 mg/mL gels display a circular morphology compared to the bipolar morphology developed by fibroblasts in 1.0-2.0 mg/mL gels (Figure 5D). Notably, matrix porosity plateaus at a minimum value at 3 mg/mL [63], likely contributing to impaired fibroblast elongation and contractility at this concentration. Therefore, it is not surprising that fibroblasts cultured in 3 mg/mL experience impaired ability to elongate, contract, and establish tensional homeostasis. Using our platform, we found that tissue-level forces in 3.0 mg/mL gels were delayed relative to the other groups and had not achieved a steady-state value by 48 hours (Figure 3B). This correlated with reduced tissue compaction (Figure 3C) and significantly lower Cauchy stress (Figure 3D-E). Transcriptional analyses at 24 hours confirmed that fibroblasts in 3.0 mg/mL displayed major changes in gene expression, including reduced proliferation [15,28,64] (Figure S15A). Reduced α-SMA expression was confirmed by lower expression of the *Acta2* gene, supporting the observed reduction in active force generation along with a mildly reduced expression of the genes *Myl9* and *Myh10* (Figure S15B-C). Interestingly, the most significantly upregulated genes in 3.0 mg/mL relative to other concentrations were *Celsr1* and *Dhcs1*, which encode proteins involved in planar cell polarity signaling [65] (Figure S15D). Upstream activation of polarity genes may indicate that fibroblasts attempt to polarize and assume an elongated morphology, which would ultimately allow them to remodel collagen through contractile forces. In terms of adhesion proteins, fibroblasts at 3.0 mg/mL maintain a fibroblastic pattern of gene expression with low *Cdh1* expression and high *Cdh2* expression (Figure S15E), along with significant upregulation of *Itga2*, the gene encoding for the α2 integrin subunit, which is key to fibroblast adhesion to collagen [66] (Figure S15E-F). Notably, fibroblasts in 3.0 mg/mL collagen do not change the expression of genes involved in ECM deposition or degradation, while they decrease the expression of ECM crosslinking genes, such as *Lox* (Figure S15G-I). This finding indicates that, on short time scales, tensional homeostasis does not involve production, degradation, or chemical modification of the ECM, but rather its reorganization via contractile forces. Finally, fibroblasts cultured in 3.0 mg/mL collagen drastically upregulated the expression of genes involved in vascular signaling – both angiogenesis and lymphangiogenesis – including *Kdr*, *Flt4*, *Vegfc*, *Gdf2*, *Sox18*, *Tek*, and *Eng*. (Figures 6E and S15J). This coordinated response may represent an effort to recruit vascular and/or lymphatic networks in response to a challenging physical microenvironment. Keesler et al. [64] previously reported upregulated mRNA expression of VEGFC by fibroblasts in anchored tissue equivalents compared to fibroblasts in free-floating tissue equivalents. However, here we suggest that upregulation of the genes encoding VEGFC (*Vegfc*) as well as its receptors VEGFR2 (*Kdr*) and VEGFR3 (*Flt4*) may underlie an autocrine signaling loop that fibroblasts activate to promote survival under conditions of altered tensional homeostasis (Figures 6F and 7).

We note that this study has a few limitations. First, we used only NIH/3T3 fibroblasts and cultured them for 48 hours. NIH/3T3 fibroblasts are widely used to study cell-ECM interactions because they can be cultured easily and transfected with high efficiency. However, they display a moderately reduced contractile ability with respect to primary (non-transformed) cells [27]. In addition to including primary fibroblasts, future studies will need to ensure that our platform is capable of culturing tissue equivalents well beyond 48 hours, to track the temporal evolution of candidate homeostatic variables. Moreover, here we cultured tissue equivalents primarily under a uniaxial constraint, rather than evaluating the proposed homeostatic metric in both uniaxially and biaxially constrained tissues. Another limitation is that our study did not utilize external perturbations in the form of either static or dynamic stretching, which would facilitate the identification of alternative metrics for the homeostatic set-point. Future incorporation of these features will facilitate mechanistic investigations on the mechanobiology of tensional homeostasis.

## 5. CONCLUSION

Our integrated biomechanical bioreactor represents a promising platform to explore tensional homeostasis in both healthy and diseased soft tissues. Beyond measuring tissue-level forces, our design integrates confocal microscopy to enable quantification of the evolving tissue microstructure and geometry. Using this platform, we found that fibroblasts do not maintain constant stress across varying collagen concentrations. Instead, tensional homeostasis may emerge from a dynamic balance between cell contractility and collagen densification. This work thus establishes a novel platform and framework for investigating the mechanobiological basis of normal and disrupted tensional homeostasis.

## Acknowledgments

We thank Adil Khan for adapting existing cell culture protocols in collagen and Bishant Karki for conducting cell density quantifications. We gratefully acknowledge Professor Jay D. Humphrey (Yale University) and Professor Patrick W. Alford (University of Minnesota) for kindly sharing prior designs of their custom devices. We thank Professor Victor D. Varner (University of Texas at Dallas) for assisting with retraction experiments and Professor W. Matthew Petroll (University of Texas Southwestern Medical Center) for fruitful conversations on cell extraction for RNA sequencing. We thank Professor Guy Genin (Washington University in St. Louis) for helpful discussions on power-law scaling. We also thank Dr. Prithvi Raj, Chaoying Liang, and Carlos Arana (University of Texas Southwestern Medical Center) for expert assistance in sequencing RNA and in conducting initial bioinformatic analyses.

## CRediT authorship contribution statement

Conceptualization: JF

Data Curation: AVG, VVN Formal Analysis: AVG, VVN Funding Acquisition: JF, GO Investigation: AVG, VVN

Methodology: AVG, DP, MDM, HH, CJC Project Administration: JF

Resources: JF

Software: AVG, VVN Supervision: JF Validation: AVG, VVN

Visualization: AVG, VVN, JF

Writing – original draft: AVG, VVN, MDM, HH, GO, CJC, JF Writing - review & editing: AVG, VVN, JF

## Data availability statement

All data in the main text or in the supplementary materials are available upon reasonable request from the corresponding author. Bioreactor design files for fabrication and custom image analysis and mechanical testing codes are openly available on GitHub https://github.com/TMR-Lab-UTD/biaxial-bioreactor. RNA-seq data will be deposited in the NCBI Gene Expression Omnibus (GEO) upon acceptance of the manuscript for publication. Data acquisition and analysis codes are also available from the corresponding authors upon reasonable request.

## Supplementary materials

Supplementary information accompany this manuscript.

