## Supplementary Material for "Dissecting the Dynamic Evolution of Tensional Homeostasis in Fibroblasts using an Integrated Biomechanical Bioreactor Platform"

### SUPPLEMENTARY METHODS

***Bioreactor pods.*** Simplified pods were designed to enable high-throughput endpoint experiments without force sensing capabilities, including tissue retraction assays, immunofluorescence staining, and RNA sequencing. A design rendering of a bioreactor pod is shown in Figure S2. Pods were 3D printed using the biocompatible resin described in the "*Biocompatible parts*" section of the manuscript. Similar to the bioreactor chamber, the bottom of the pod is open to accommodate the 100 mm petri dish, while the pod lid has a 35 mm glass coverslip bonded at the center to facilitate imaging. Bioreactor pods were cultured in a standard cell culture incubator for an amount of time that depended on the experimental endpoint. To facilitate confocal and multiphoton imaging, pods were secured to dedicated baseplates. Each pod was reused (after autoclaving) for up to 16 experiments before disposal.

***Porous Hydrogel InserTs (PHIT).*** Several methods have been proposed in the literature to transmit loads from the arms to the collagen gels, including embedded polypropylene mesh [1], porous polyethylene bars [2], and sutures [3]. Here, we utilized 3D printed PHIT by adapting the design of the porous inserts previously introduced by Eichinger et al. [4]. There are separate PHIT variants for mounting tissues on the passive side and on the force sensing side of each axis (Figure S4). In particular, the PHIT for the force transducer is mounted using a 29-gauge wire which is inserted into the PHIT between the drying and curing process, using an additional small amount of resin as adhesive. The wire is then inserted into the probe of the force transducer and secured with Crytsalbond wax (SPI Supplies, Crystalbond 509) in combination with a custom jig for precise PHIT alignment. Without such jig, the PHIT may bend over time under the action of gravity, contacting the bottom of the petri dish, and introducing severe noise into the force data. The PHIT were 3D printed using a biocompatible resin, as described in the manuscript. Once mounted to the force transducers, the PHIT were sterilized in a 10% bleach solution for 5 minutes, washed out with $dH_2O$, and immersed in 70% ethanol until further assembly was ready. A 100 mm petri dish with tissue molds (prepared as described in the section "*Tissue molds*") was inserted into the chamber to accommodate the PHIT. Excess ethanol was aspirated from the PHIT prior to mounting on the device so that it did not seep under the molds and break the seal with the petri dish.

***Tissue molds.*** Polydimethylsiloxane (PDMS) molds to form collagen gels in a dogbone (uniaxial configuration) or cruciform (biaxial configuration) shape were prepared using the Sylgard 184 Silicone Elastomer Kit (Krayden, DC4019862). Images of the dogbone and cruciform molds are shown in Figure S5. The silicone elastomer base (Part A) was mixed with the silicone elastomer curing agent (Part B) in a 10:1 w/w ratio. A mass of 17-to-20 grams of the mix was poured into an inverse mold made of a thermoplastic polyurethane filament (TPU), shore hardness 95A (SainSmart, thermoplastic polyurethane) In turn, the TPU inverse mold was 3D printed using a fused deposition modeling (FDM) 3D printer (Bambu Lab, P1P), with a 0.2 mm layer height, 2 perimeters, 15% cross hatch infill, and ironing enabled on top surfaces. Once poured, the PDMS mold was placed into a vacuum-sealed oven for 60 minutes and then heated at 55°C overnight. PDMS molds were cut into quarters, sterilized in 70% ethanol under UV light for 15 minutes, and air-dried overnight. The molds were aligned in a 100 mm plastic petri dish and filled with 3 mL (uniaxial configuration) or 6 mL (biaxial configuration) of a 10 mg/mL sterile bovine serum albumin (BSA) solution, which was used to passivate the petri dish for 30 minutes and then aspirated. The dish was added to either the main biomechanical bioreactor or a bioreactor pod and preheated in a 37ºC incubator for 1 hour.

***Collagen remodeling.*** We analyzed local changes in collagen microstructure by processing the Best-Z slice from the CRM channel. The Best-Z slice is defined as the maximum intensity projection of the substack obtained by selecting three images above and three images below the manually tracked bottom surface of the gel. Therefore, the Best-Z slice represents a time series of images capturing the structural evolution at the gel bottom, where the image contrast is highest. The corresponding TD time series was selected from the middle slice of the substack tracking the bottom surface of the gel. The area of the collagen gel in the Best-Z slice was determined using an Otsu threshold on the TD channel for each time frame. The CRM intensity was calculated from the Best-Z slice as the average pixel value within the segmented area (i.e., area occupied by the collagen gel). The temporal evolution of the CRM intensity was then used to estimate temporal changes in collagen density. The fiber organization was analyzed using the ImageJ plugin Ridge Detection [5] within the entire image. Images were converted into an 8-bit format and analyzed using the following settings for Ridge Detection: line_width = 20, low_contrast = 30, high_contrast = 160, min_length = 20, max_length = 0, dark_line = False, estimate_width = True, extend_line = True, correct_pos = False. Ridge Detection produces a list of fiber locations and angles which were output to a CSV file. Fibers in the 181°-360° range were remapped to the 0°-180° range. The percentage of vertical fibers (parallel to the principal axis of the bioreactor) was calculated as the fraction of fibers with angles in the 85°-95° range. To quantify collagen alignment dynamics, a logistic function $y(t) = A/[1 + \exp((t_0 - t)/\tau\ )\ ] + B$ was fitted to the percentage of vertical fibers plotted over time for each collagen concentration, where $A$ represents the difference between the percentage of vertical fibers at steady-state and initial state, $B$ represents the percentage of vertical fibers at the initial state, $t_0$ is the time at the inflection point, and $\tau$ represents the alignment time constant.

***Cross-sectional area estimation.*** A custom Python pipeline was developed to measure the cross-sectional area changes in the gel during compaction (Figure S9A). For each time point, the CRM channel in the tiled z-stack was resliced to visualize the cross-section of the gel. To circumvent the limited penetration depth of CRM imaging, we developed a circular arc extrapolation method to estimate the top surface geometry of the gel. First, we identified the bottom edge of the gel as the lowest pixel in each column of the image above an automatically determined Otsu threshold. Second, the four original corners of the gel ($a$, $b$, $c$, and $d$, numbered counterclockwise starting from the top left) were manually identified for each frame of the resliced time series. Third, a 3-point arc was drawn from point $a$ to $d$ with center point $P$, where $P = \frac{1}{2}(P_{ab} + P_{dc})$, where $P_{ab}$ is the intersection of $ab$ and the perpendicular bisector of $ad$, and $P_{dc}$ is the intersection of $dc$ and the perpendicular bisector of $ad$. The arc was drawn using the shorter path (clockwise or counterclockwise) if both points $a$ and $d$ lie above line $bc$, otherwise the arc was drawn clockwise from point $a$ to $d$. The cross-sectional area was calculated as the area of the polygon bounded by the segmented bottom edge and the extrapolated top surface (Figure S9B).

***Multiphoton microscopy.*** We imaged the central region of fresh tissue equivalents using a multiphoton microscope (Olympus, FV MPE-RS) at the UT Dallas Imaging & Histology Core. Two laser beams (Spectra Physics, Insight DS+ and Mai-Tai HP DeepSee) were focused on the tissues through a 10× water-immersion objective (Olympus, 0.6 N.A., 8 mm working distance) mounted in upright configuration. One laser beam was set to an excitation wavelength of 1060 nm (Insight DS+) and Second Harmonic Generation (SHG) signal from the collagen matrix was collected using a 495-540 nm bandpass filter, while the second laser beam was set to an excitation wavelength of 740 nm (Mai-Tai HP DeepSee) and the autofluorescence signal from cellular nicotinamide adenine dinucleotide (NADH) was collected using a 460-500 nm bandpass filter.

Acquisition of tiled image stacks used the following settings: 1024 × 1024 pixels at a resolution of 1.243 μm/pixel, a pixel dwell time of 4 μs, a stack size of 3,900 μm with 50 μm steps, and a 10% overlap. To obtain the ground-truth cross-sectional area for our tissue equivalents, we prepared multiple uniaxial dogbone samples in the bioreactor pods using a collagen concentration of 0.8 mg/mL and a cell density of 500,000 cells/mL, cultured them for varying time frames (0, 4, 6, 9, 12, 18, 24, and 48 hours), and imaged them using both confocal and multiphoton imaging modalities. The cross-sectional areas from confocal microscopy were estimated as described in the supplementary methods. Instead, cross-sectional areas from multiphoton microscopy were estimated by reslicing the SHG/NADH channels in the tiled z-stack and manually tracing the cross-sectional area of the collagen gel occupied by fibroblasts. To validate our cross-sectional area estimation approach, we employed linear regression to assess how closely the "Estimated Area" obtained from confocal microscopy matched the ground-truth "MPM Area".

We also used multiphoton imaging to quantify the cell density within fixed tissue equivalents. Samples used for confocal time-lapse imaging with synchronous force measurement for 48 hours were detached from the PHIT using fine scissors and fixed overnight using cold 4% paraformaldehyde (PFA, Fisher, AAJ19943K2). After washing twice with 1× Phosphate Buffered Saline (PBS), the fixed tissue equivalents were optically cleared using CUBIC [6] (Clear, Unobstructed Brain/Body Imaging Cocktails) following established protocols [7]. We prepared a mixture of 25 wt% urea (Fisher, U15), 25 wt% Quadrol (N,N,N',N'-Tetrakis(2-hydroxypropyl)ethylenediamine, Sigma-Aldrich, 122262), 15 wt% Triton X-100 (Sigma-Aldrich, T8787), and $dH_2O$. Samples were pretreated for 3 hours in 4 mL of 50 vol% diluted CUBIC reagent (1:1 with $dH_2O$) and then immersed in 2.5 mL CUBIC reagent under gentle shaking at room temperature. The CUBIC reagent was refreshed every two days and samples were cleared for 25 to 31 days. To ensure proper cell staining across the thickness of the gel, 5.71 µL of DAPI (Invitrogen, D1306, 1:1000) was added for 7 days prior to imaging. The 3D organization of optically cleared tissue equivalents was investigated using multiphoton microscopy. The laser beam was set to an excitation wavelength of 1060 nm (Insight DS+) and a 10× water-immersion objective (Olympus, 0.6 N.A., 8 mm working distance) was used to focus the excitation light on the central region of the tissue equivalents. We collected the DAPI signal from fibroblasts using a 410-455 nm bandpass filter and the SHG signal from the collagen matrix using a 495-540 nm bandpass filter. Acquisition of tiled image stacks used the following settings: 1024 × 1024 pixels at a resolution of 1.243 µm/pixel, a pixel dwell time of 4 µs, a stack size of 1900 µm with 10 µm steps, and a 10% overlap. The resulting 3D tiled stacks were analyzed using Imaris v10.1.1 (Oxford Instruments). DAPI-stained nuclei were segmented using spot detection with an estimated XY diameter of 18 µm and Z diameter of 45 µm, with background subtraction enabled. Detected spots were filtered by quality (threshold > 5.0) and classified using machine learning to distinguish true nuclei from artifacts. The SHG signal from collagen was used to determine the volume of the compacted tissue using surface segmentation with a surface grain size of 2.49 µm and smoothing enabled. Surfaces were filtered to retain only those with >10 voxels. Cell density was calculated as the number of classified DAPI-positive spots divided by the compacted tissue volume.

***Viability assay.*** Cell viability was assessed after 48 hours of culture using a live/dead staining kit (Invitrogen, L3224). The experimental media in the bioreactor was aspirated and replaced with 25 mL of culture media (DMEM supplemented with 10% FBS and 1% Pen/Strep) containing 25 µL ethidium homodimer-1 (EthD-1) and 6.67 µL calcein AM. The bioreactor was incubated at 37°C and 5% $CO_2$ for one hour, after which the staining media was aspirated and replaced with fresh culture media. Confocal imaging was performed using a 10× objective with a field of view of 1024 × 1024 pixels at a resolution of 0.752 μm/pixel and a pixel dwell time of 0.5 μs. Image stacks were acquired using 1,750 μm z-stacks with a 50 μm step size. Tiling was employed to capture the entire gel width using 5 image tiles with 10% overlap. Two fluorescence channels were acquired: calcein AM (live cells, excitation 488 nm / emission 500-530 nm), EthD-1 (dead cells, excitation 561 nm / emission 570-616 nm). Resliced z-stacks were used to visualize the distribution of live and dead cells (Figure S7).

***Bulk RNA sequencing.*** Transcriptome-wide gene expression of NIH/3T3 fibroblasts was analyzed using bulk RNA-sequencing (RNA-seq). Steps in the experimental and analysis pipeline are described below.

*Sample preparation.* For each of the four collagen concentrations (1.0, 1.5, 2.0, and 3.0 mg/mL), three replicates – each at a different cell passage – were cultured in pods for 24 hours. Subsequently, cells were isolated from the gel by cutting the central region of the gel between the porous hydrogel inserts and incubating it in 2 mg/mL collagenase (Gibco, 17100017) in DMEM supplemented with 1% Pen/Strep. The gels were digested at 37°C for 45 minutes with gentle mixing using a HulaMixer Sample Mixer (Fisher, 15920D). Cells were pelleted by centrifugation at 1500 RPM for 4 minutes, and the supernatant was aspirated. The cell pellet was resuspended in 1 mL of TRIzol (Invitrogen, 15596026), homogenized by passing the cell pellet through a 20-gauge needle 10 times, and stored at -80°C until RNA extraction.

*RNA extraction and sequencing.* RNA extraction was performed using an RNeasy Mini Kit (Qiagen, 74104) and TRIzol Reagent, following the manufacturers' instructions. RNA quality was assessed using the Agilent Bioanalyzer with the RNA Nano Chips Kit (Agilent, 5067-1511). Concentration was measured with a spectrophotometer (Thermo Scientific, NanoDrop 2000). The RNA Integrity Number (RIN) value for all 12 selected RNA samples was greater than 7. Libraries were prepared using the KAPA RNA HyperPrep Kit with RiboErase (HMR) (Roche, KK8561), according to the manufacturer's instructions. A total of 12 RNA libraries were constructed using 2.5 µg of RNA input per sample. The workflow included: rRNA depletion, RNAse H and DNase treatment, fragmentation, first-strand cDNA synthesis using reverse transcriptase and random primers, second-strand synthesis and A-tailing, with dUTP incorporated into the second strand, adapter ligation using UMI adapters synthesized by IDT (Integrated DNA Technologies), PCR enrichment with 10 cycles. Final libraries were purified using AMPure XP beads, quantified using PicoGreen, and assessed for quality on the Agilent Bioanalyzer using the High Sensitivity DNA Kit (Agilent, 5067-4626). Libraries were sequenced on an Illumina NovaSeq X platform using a PE-110 protocol. Each sample yielded approximately 40 million passing filter reads.

*Differential expression analysis.* Paired end demultiplexed fastq files were generated using bcl2fastq2 (Illumina, v2.20), from NovaSeq 6000 bcl files. Initial quality control was performed using FastQC v0.11.8 and multiqc v1.7. Fastq files were imported batch wise, trimmed for adapter sequences followed by quality trimming using CLC Genomics Workbench (CLC Bio, v24.0.2). The imported high-quality reads were mapped against gene regions and transcripts annotated by ENSEMBL v110 GRCm39 using the RNA-Seq Analysis tool v2.8 (CLC Genomics Workbench), only matches to the reverse strand of the genes were accepted (using the strand-specific reverse option). The resulting RNA expression browser was filtered for protein coding genes which have a raw count of at least 10 for all replicates of any of the initial concentrations. A principal component analysis (PCA) plot was generated from the raw counts after applying a variance stabilizing transform (VST) provided by the PyDeseq2 library [8]. Differential expression analysis was performed using PyDeseq2 for 1.5, 2.0, and 3.0mg/mL gels with the 1.0 mg/mL gels as the reference (n = 3 biological replicates/group). Statistically significant genes were considered as those with a p-value < 0.05 and $\log_2$-fold-change > $\log_2(1.5)$. Pathway overrepresentation analysis was performed using GSEApy [9] and Enrichr [10] with the "Reactome_Pathways_2024" database [11]. Enrichr overrepresentation was run separately using the list of significantly upregulated DEGs and downregulated DEGs for each comparison without specifying a list of background genes.

### SUPPLEMENTARY DISCUSSION

Over the course of more than four decades, numerous studies have used tissue equivalents to examine the mechanisms by which fibroblasts remodel 3D collagen networks through the generation of endogenous contractile forces. Early on, the existence of contractile forces was revealed by the dramatic compaction observed in free-floating tissue equivalents. In a pioneering paper, Bell et al. [12] showed that fibroblasts compact the collagen network in the absence of external constraints. The final degree of compaction was determined by the initial collagen concentration $\rho_0$ as well as by the initial cell density, among other factors [12,13]. Subsequent studies revealed that such macroscopic compaction results in a net increase in the final collagen density [13,14]. Consistent with our observations, the microscopic densification of collagen in free-floating tissue equivalents depends on the initial collagen concentration $\rho_0$: tissue equivalents with a lower $\rho_0$ become denser over time than tissue equivalents with a higher $\rho_0$, even though the total collagen mass remains lower than in initially denser tissues [14]. This extreme collagen densification results from the highly compressible behavior of collagen networks [15], which, starting from an initially isotropic microstructure undergo heterogeneous remodeling in response to anisotropic cell contraction [16], leading to the development of a residually stressed structure with compression at its center and tension at its periphery [17]. The residual stress field is created by fibroblasts as they generate a tensile ring of densely aligned cells and collagen [17]. Contraction of this tensile ring compacts the gel by squeezing out the interstitial fluid. Therefore, contrary to the commonly held assumption that free-floating tissue equivalents are mechanically relaxed [18,19], tension-driven tissue remodeling drives gel compaction as fibroblasts attempt to achieve tensional homeostasis.

Anchoring tissue equivalents to the dish mechanically restrains one of the gel surfaces [18]. Remodeling of these anchored tissue equivalents over time can be measured macroscopically by tracking the reduction in gel thickness [20] and microscopically by monitoring the acquisition of an elongated cell morphology [21]. Under these conditions, fibroblasts acquire a myofibroblast phenotype with visible stress fibers that disappear rapidly after the release of mechanical tension [22]. Indeed, mechanical tension in fibroblasts has been associated with a proliferative and synthetic phenotype [23]. Conversely, release of mechanical tension triggers an apoptotic response in fibroblasts [24,25]. While deeply insightful, studies involving free-floating and anchored tissue equivalents were limited by the inability to measure endogenous forces and to impose exogenous loads. In the 1990s, R.A. Brown began to fill these technological gaps by developing the culture force monitor (CFM) [26] and the tensioning culture force monitor (t-CFM) [27]. Much like previous isometric force monitors [28–30], both CFM and t-CFM were designed to constrain tissue equivalents at two opposite boundaries while measuring contractile forces via a transducer connected to one boundary. Similar to our tissue-level force data (Figure 1), the tissue-level forces measured by the CFM followed a sigmoidal time-course and were associated with the initiation and extension of filopodial processes into the collagen matrix [31]. More importantly, the simultaneous application and measurement of forces via the t-CFM revealed the existence of a preferred level of mechanical tension, which led to the introduction of the concept of *tensional homeostasis* [27]. Additionally, the t-CFM facilitated the discovery that a remodeled collagen network bears a significant portion of the homeostatic tension [32]. Studies that employed a similar exogenous loading platform [33] identified that actin integrity, myosin activation, and binding to collagen via the β1 integrin subunit play a key role in the development of homeostatic tension [34].

While these studies clearly indicated that tensional homeostasis emerges from a feedback loop between cell contractility and ECM remodeling, they did not reproduce the biaxial loading state experienced by many soft tissues in vivo [35]. Thus, biaxial bioreactors were proposed as versatile systems to analyze tissue equivalents under the action of semi-physiologic loads [36]. These platforms offered some key advantages over the CFM/t-CFM technology. First, they included biomechanical testing protocols to explore how fibroblasts develop mechanical anisotropy in tissue equivalents [37]. Second, they enabled modulation of the degree of anisotropy by using uniaxial, biaxial, or intermediate (strip biaxial) loading modalities [3]. Third, they integrated optical access for advanced microscopy techniques to monitor collagen organization, such as multiphoton SHG, that were used to investigate the structural remodeling of collagen by stromal fibroblasts [2,38,39]. Among other findings, these systems revealed that collagen is deposited and aligned along the direction of principal stress while the tissue equivalent compacts by losing volume due to compressibilty [38]. Importantly, the bioreactor developed by Eichinger et al. [4] was used to show that homeostatic states can be achieved by tissue equivalents under uniaxial, biaxial, or strip biaxial constraints [3]. This indicates that fibroblasts establish tensional homeostasis regardless of the mechanical constraint they are exposed to. Additionally, Eichinger et al. [4] showed that the homeostatic tension increases quasi-linearly as a function of cell density and initial collagen concentration $\rho_0$, confirming earlier observations made by Delvoye et al. [29]. The fact that the homeostatic tension is not preserved in various experimental conditions suggests that the tissue-level force is not preserved by fibroblasts, a conclusion that was also reached via theoretical and computational modeling [40]. Instead, the homeostatic set-point is more likely to be related to microscale structural or mechanical variables [41], which are naturally assessed using microfabricated systems. One such system was introduced by Legant et al. [42] and consist of silicon microcantilevers that constrain the remodeling of collagen by cells either uniaxially or biaxially. More recently, Walker et al. [43] introduced uniaxial stretching via vacuum-driven actuation and showed that fibroblasts maintain a nearly constant mean tension across strain amplitudes, therefore providing evidence for tensional homeostasis at the microscale. Microfabricated systems have also been adapted to examine the existence of tensional homeostasis in single fibroblasts using an atomic force microscopy (AFM) based assay [44]. This system revealed that the concept of tensional homeostasis does not apply at the single cell level, because single cells respond to externally applied strains by minimizing their internal tension and accommodating deformations, a behavior that has been called *tensional buffering* [44]. Taken together, results from a large body of work in this research field indicate that tensional homeostasis is an emergent phenomenon arising from mechanobiological interactions between fibroblasts and extracellular collagen.

### SUPPLEMENTARY FIGURES

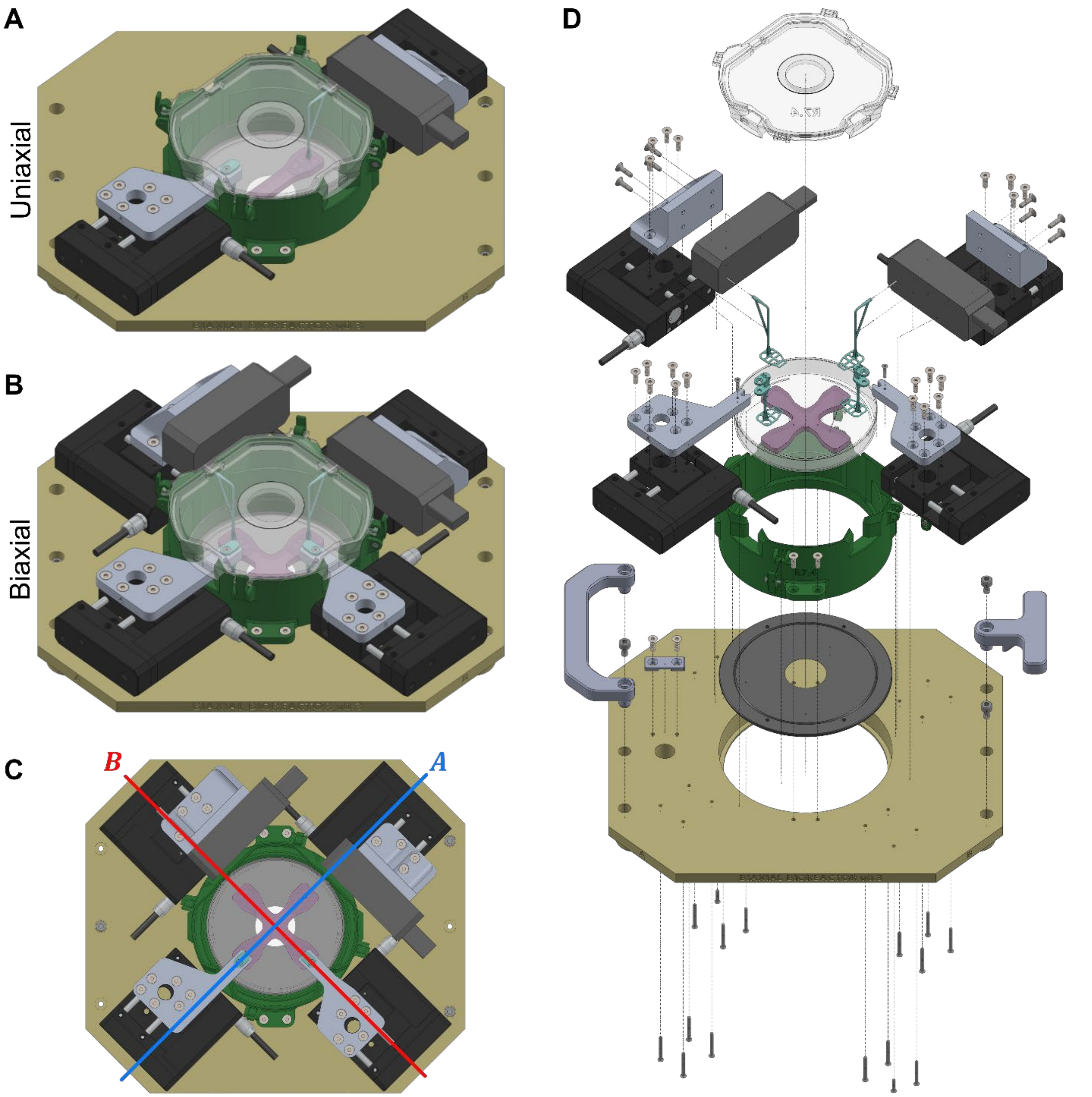


***Figure S1: Main bioreactor CAD*** Computer aided design (CAD) rendering of the biomechanical bioreactor in SolidWorks in **(A)** uniaxial and **(B)** biaxial configurations. **(C)** The main axes of the bioreactor are oriented at 45° relative to the baseplate to accommodate the geometrical constraints of the confocal microscope stage. (**D**) An exploded view of the biaxial configuration to highlight individual components.

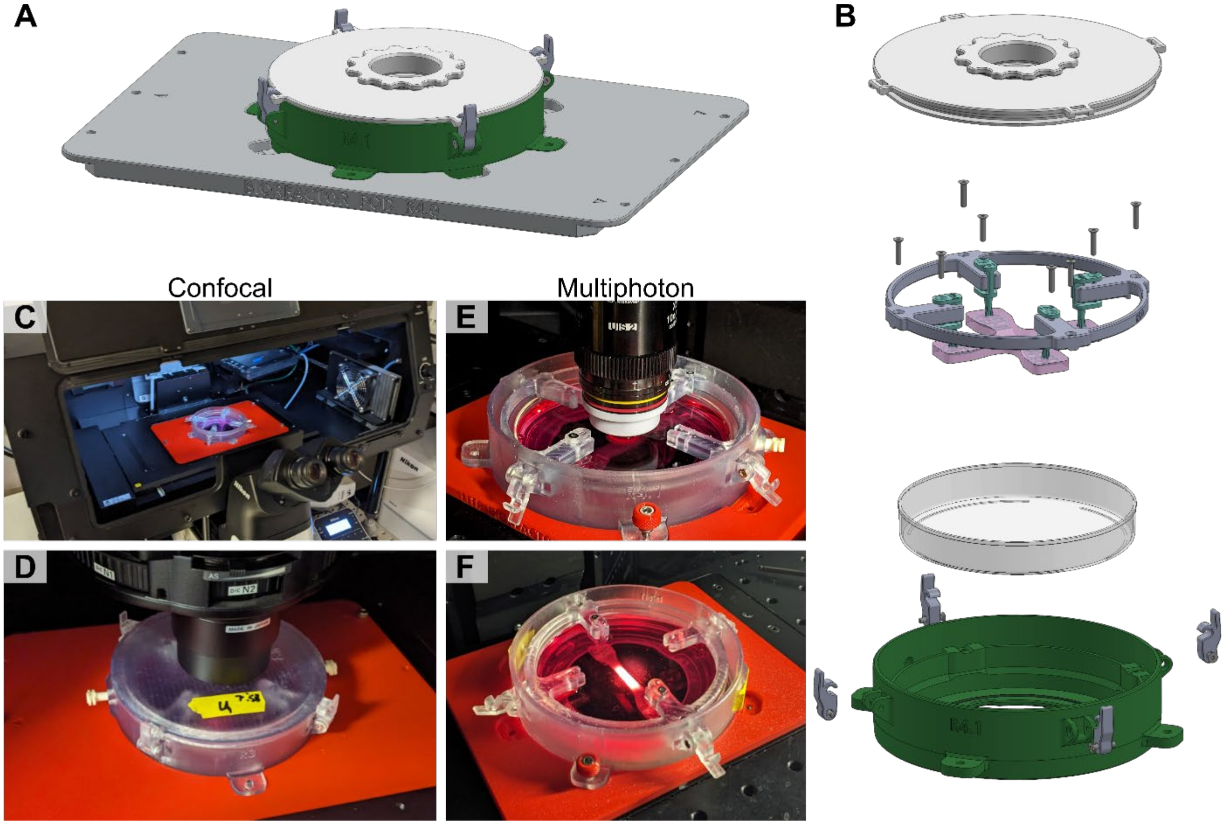


***Figure S2: Bioreactor pod CAD*** **(A)** Computer aided design rendering in SolidWorks of a bioreactor pod fixed to a custom mount for a confocal or multiphoton microscope. **(B)** Exploded view of the bioreactor pod to highlight individual components. **(C)** A bioreactor pod mounted on a confocal microscope with the lid open. **(D)** A closer view of a bioreactor pod mounted on a confocal microscope with the lid closed and the condenser lowered. **(E)** Using a different adapter plate, the same bioreactor pod is mounted on a multiphoton microscope with a water immersion objective and **(F)** without the objective to show the uniaxially constrained tissue mounted within.

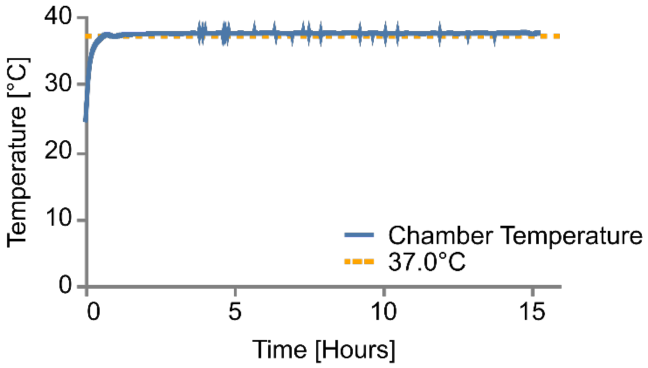


***Figure S3: Chamber temperature*** To determine the necessary settings for the ThermoPlate, we ran a test with a petri dish containing 25mL of phosphate buffered saline (PBS) inside the chamber and ThermoBox. A temperature sensor paired with the ThermoPlate was taped down to the bottom of the dish prior to filling it with 1× PBS. The internal temperature of the ThermoPlate as well as the reading from the temperature sensor were logged every 15 seconds. The heater was set to 38.0°C for the full duration of the test. After reaching steady-state, the PBS in the petri dish maintained a temperature of 37.4 ±0.06°C (mean ± standard deviation), first reaching that temperature after 45 minutes.

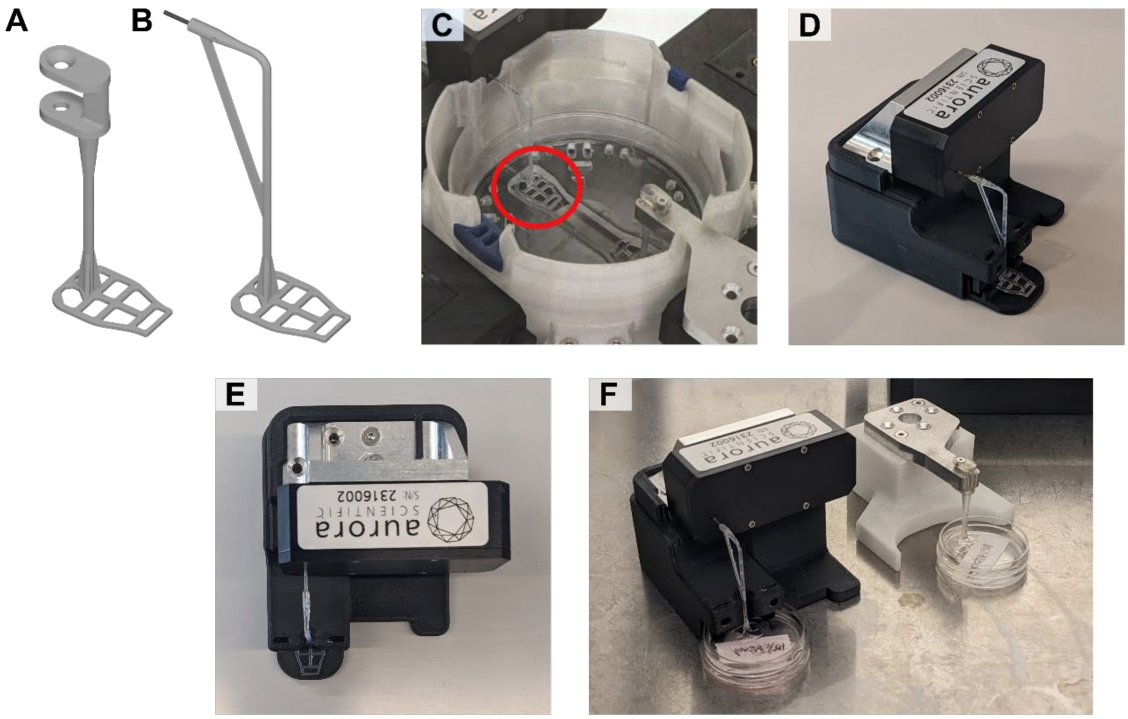


***Figure S4: Porous Hydrogel InserTs (PHIT)*** **(A)** Porous hydrogel inserts have been modified from the work of Eichinger et al. [4] to be compatible with the new platform. **(B)** Notably, due to the geometry of the force transducer and space constraints within the microscope, the shaft features a 90° bend and is printed as part of the PHIT instead of mounting via needles. **(C)** The PHIT have approximately 1 mm of clearance with the sides of the mold and bottom of the petri dish, contacting with either of which may introduce severe noise into the experiment or risk damaging the force transducer. **(D)** To precisely align the PHIT in the device, a custom mounting jig was developed. **(E)** A top view of the mounting jig with the PHIT mounted. **(F)** The mounting jig features a cutout for a 35 mm petri dish with bleach to sterilize the PHIT before mounting onto the main bioreactor.

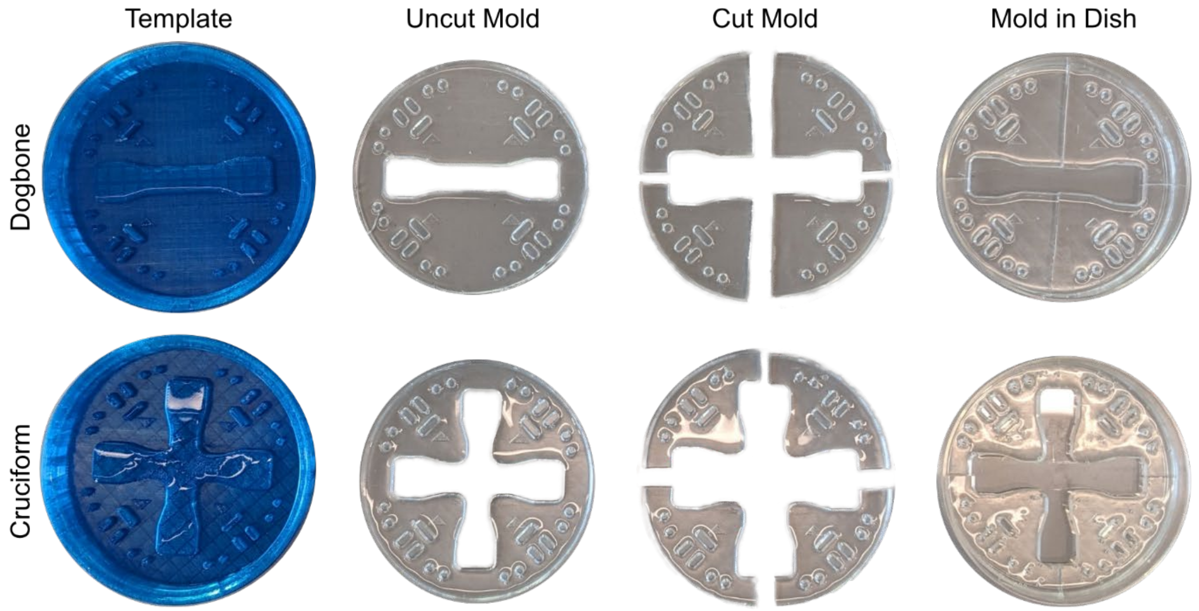


***Figure S5: Tissue molds*** Dogbone or uniaxial sample (top row). Cruciform or biaxial sample (bottom row). 3D printed TPU template for producing molds (first column). The base PDMS mold produced by the template (second column). PDMS mold cut into quarters (third column). The four quarters are finally reassembled in a petri dish (fourth column). Note the arrows on the quarters are all aligned.

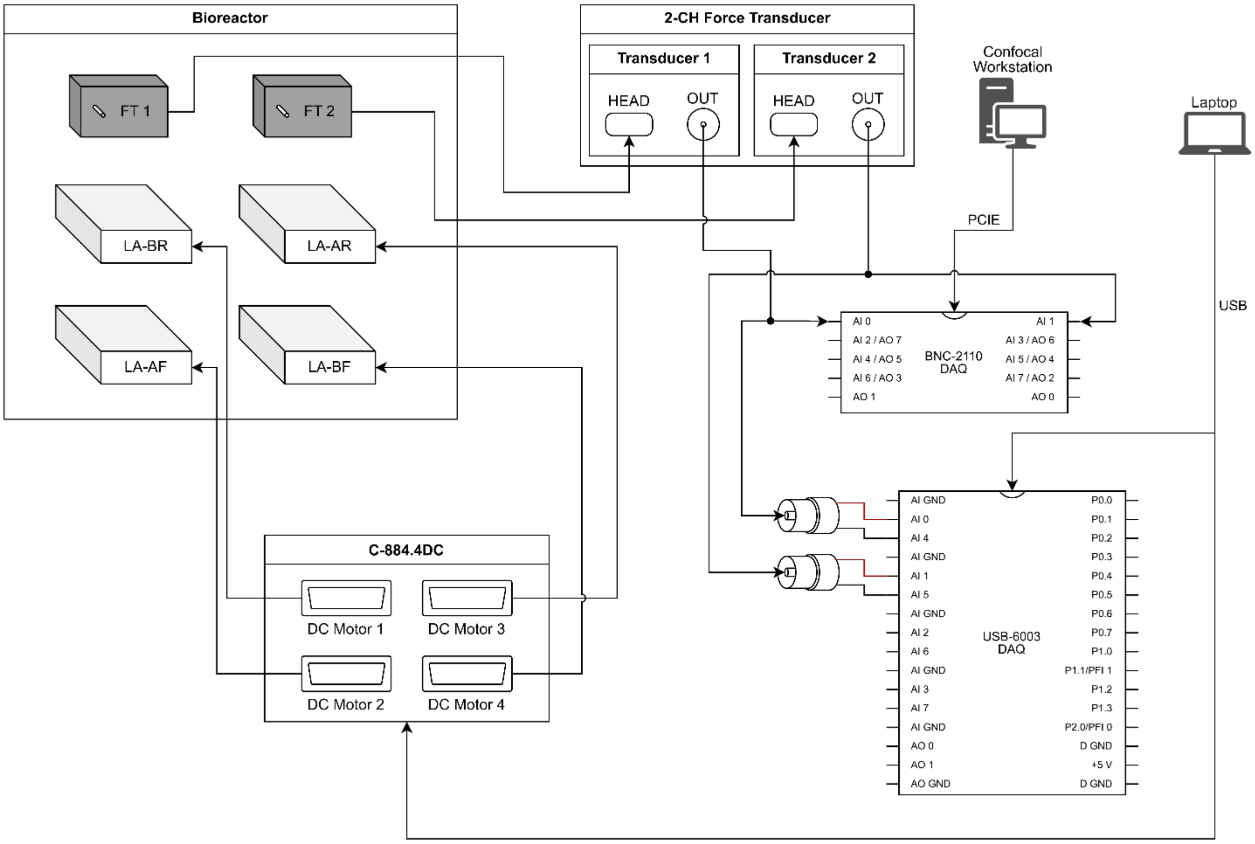


***Figure S6: Wiring schematic for mechanical stretcher*** FT = Force Transducer (Aurora Scientific, 403C); LA = Linear Actuator (Physik Insturmente, M111-1DG1). Note: The USB-6003 DAQ should be configured to operate in differential mode when wired as shown above.

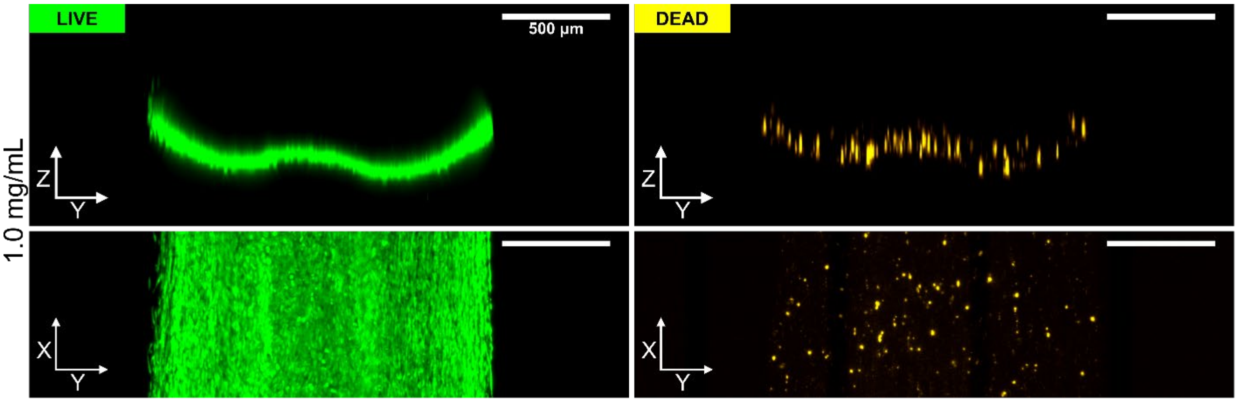


***Figure S7: Live / dead staining.*** Max intensity projection of XY and YZ of a representative 1.0 mg/mL gel after 48 hours of remodeling stained for calcein AM (live) and EthD-1 (dead).

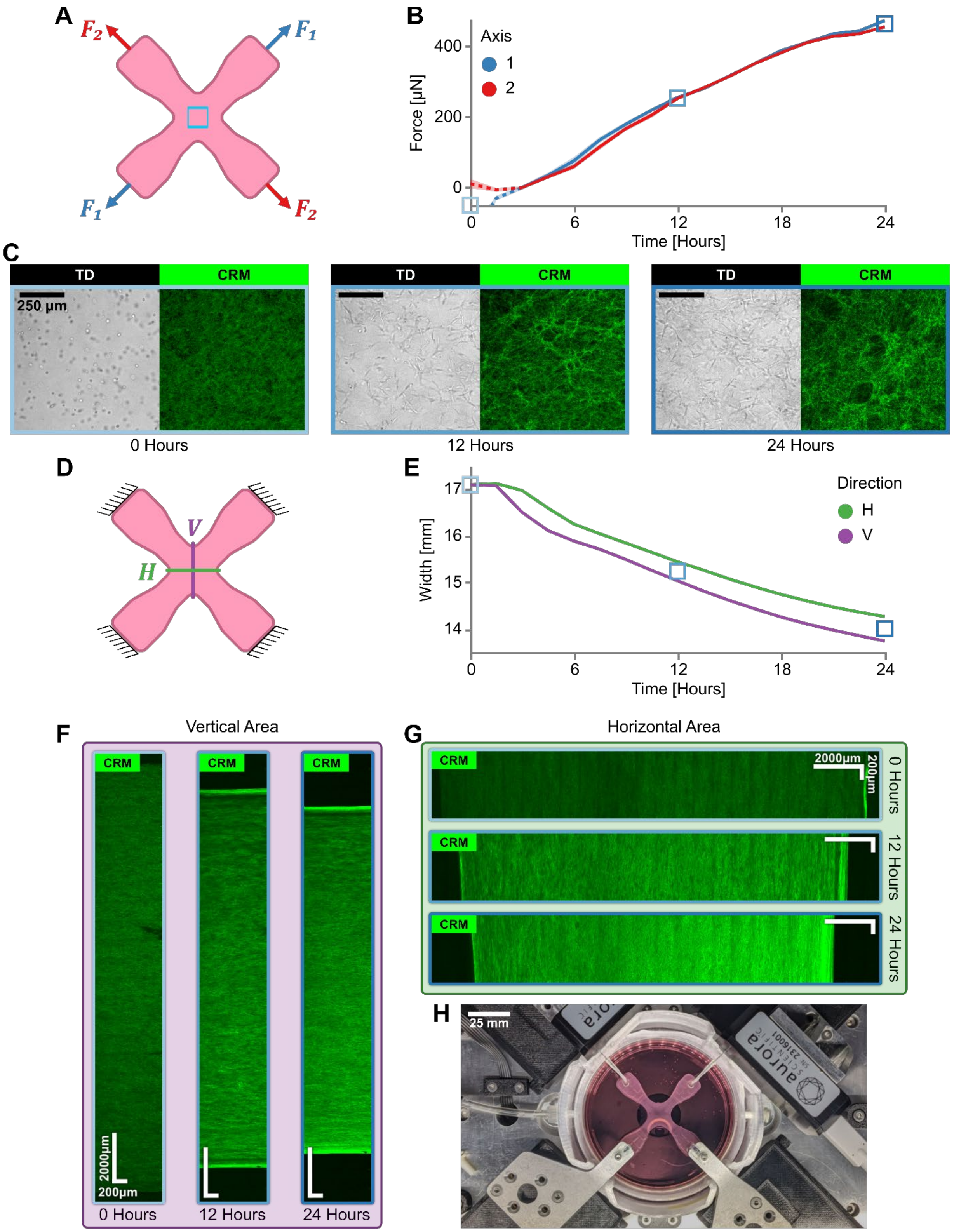
A
$F_2$
$F_1$
$F_1$
$F_2$
B
Axis
1
2
Force [µN]
400
200
0
Time [Hours]
0
6
12
18
24
C
TD
CRM
250 µm
0 Hours
12 Hours
24 Hours
D
V
H
E
Direction
H
V
Width [mm]
17
16
15
14
Time [Hours]
F
Vertical Area
CRM
2000µm
200µm
0 Hours
12 Hours
24 Hours
G
Horizontal Area
CRM
2000µm
200µm
0 Hours
12 Hours
24 Hours
H
25 mm
aurora

***Figure S8: Biaxial experiment proof of concept.*** **(A)** Free body diagram of global tension in biaxial gel along the principal axes $F_1$ and $F_2$. **(B)** Forces over 24 hours for a biaxially constrained gel in a representative sample (initial collagen concentration 1.0 mg/mL, 500,000 cells/mL). The first 3 hours are cut off due to force transducer drift transience. **(C)** Transmitted light detector (TD) and confocal reflection microscopy (CRM) images of the same gel at 0, 12, and 24 hours. Scale bars, 250 μm. **(D)** Schematic of the horizontal and vertical imaging directions, $H$ and $V$, respectively, **(E)** quantified over time. **(F)** Representative widths of the biaxial gel in the vertical axis and **(G)** horizontal axis at 0, 12, and 24 hours. Scale bar in the long axis is 2000 μm and short axis is 200 μm. The short axis is scaled up 3× the original dimension. **(H)** Overview of the system with a biaxial gel mounted after 24 hours of remodeling. Using this biaxial setup, we observed isotropic force generation accompanied by macroscopic tissue compaction and microscopic bundling of collagen fibers around randomly distributed bipolar fibroblasts.

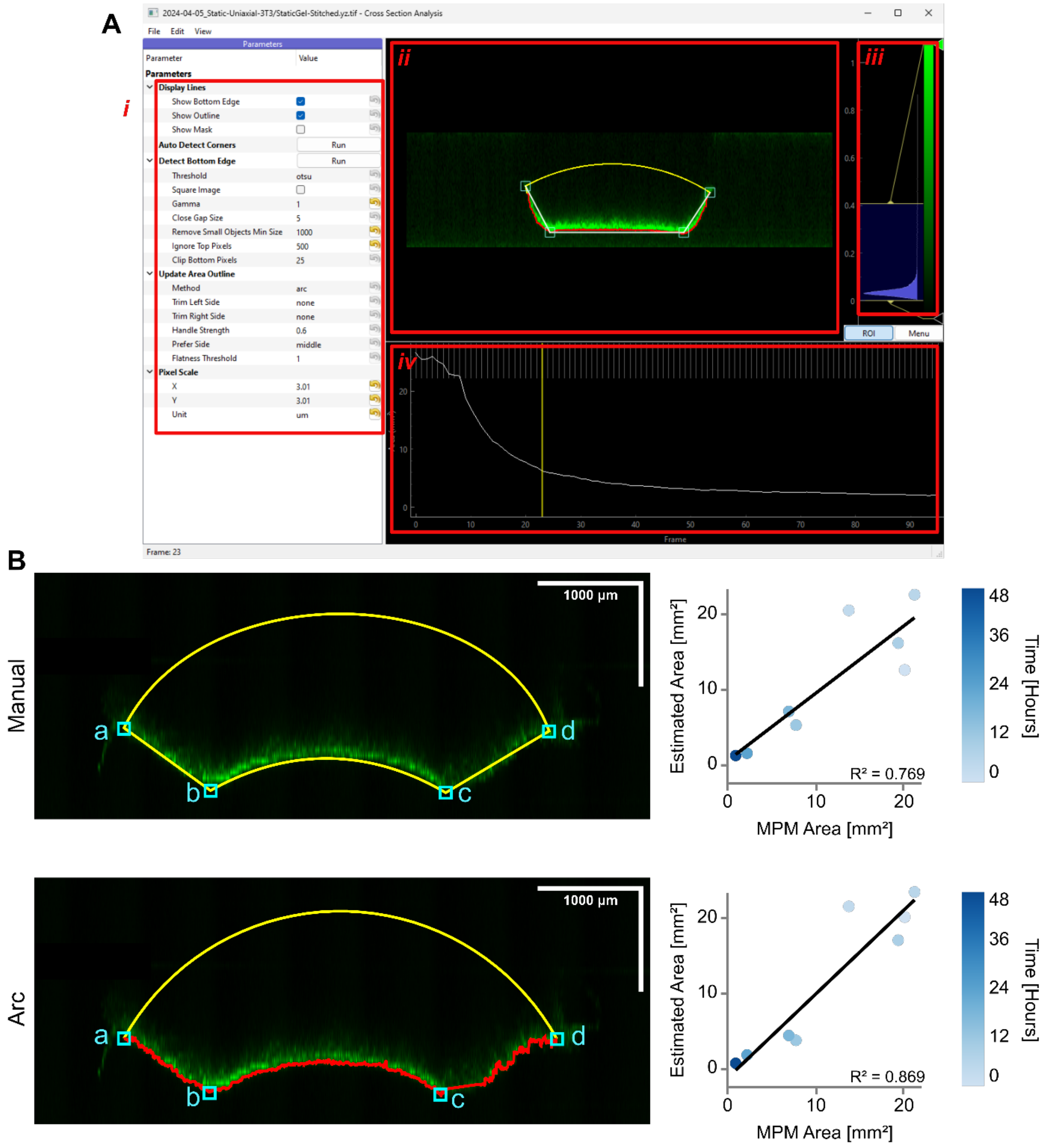


***Figure S9: Cross section extrapolation methods comparison.*** (**A**) Custom graphical user interface for processing uniaxial cross section timelapses. *i*. left panel with analysis configuration options; *ii*. Viewport showing the cross section at a given timestep with the arc cross section overlaid; *iii*. Adjustable look up table for viewing the cross section; *iv*. Timeline with cross sectional area plotted at each frame. (**B**) Comparison of manual and arc methods for extrapolating the cross-sectional area from CRM images at a representative 12-hour time point. The estimated area at each time point is plotted against the corresponding multiphoton microscopy (MPM) image of the same gel which has been manually segmented to be used as the ground truth.

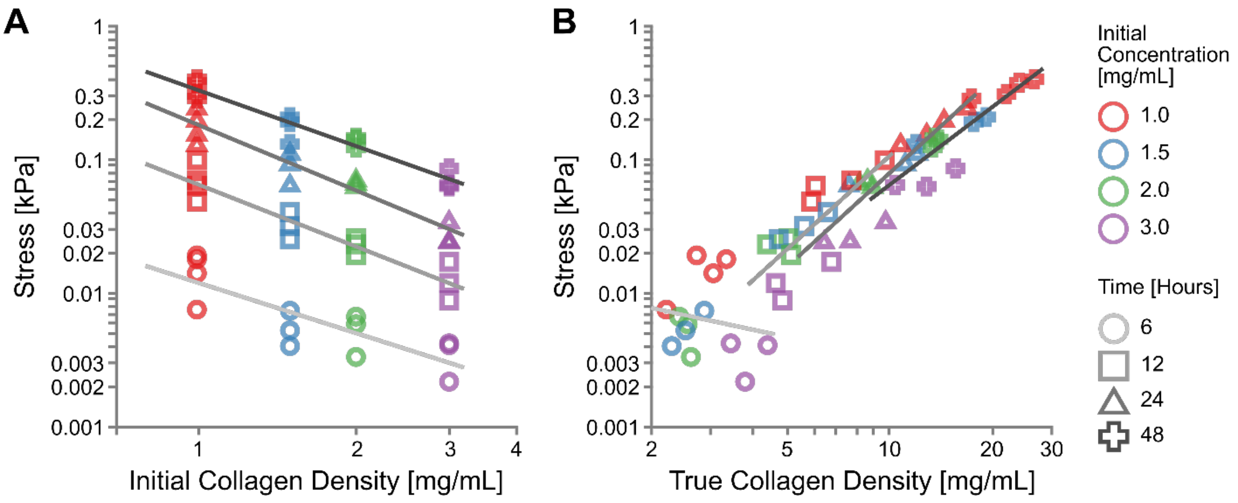


***Figure S10: Scaling relationships.*** (**A**) Inverse scaling of the Cauchy stress $\sigma$ with the initial collagen density $\rho_0$, shown at 6 hours (slope=-1.26, $R^2$=0.68, p=0.000509), 12 hours (slope=-1.56, $R^2$=0.89, p=1.36e-06), 24 hours (slope=-1.64, $R^2$=0.91, p=3.29e-07), and 48 hours (slope=-1.40, $R^2$=0.93, p=9.17e-08). (**B**) Power law scaling of the Cauchy stress $\sigma$ with the true collagen density $\rho$, shown at 6 hours (slope=-0.55, $R^2$=0.03, p=0.58), 12 hours (slope=2.32, $R^2$=0.56, p=0.00321), 24 hours (slope=2.33, $R^2$=0.79, p=5.19e-05), and 48 hours (slope=1.93, $R^2$=0.78, p=5.87e-05). Data are presented on a log-log scale. The inverse scaling remains consistent across all time points, while the sign of the slope changes from negative to positive over time and the goodness of fit of the power law relationship improves progressively.

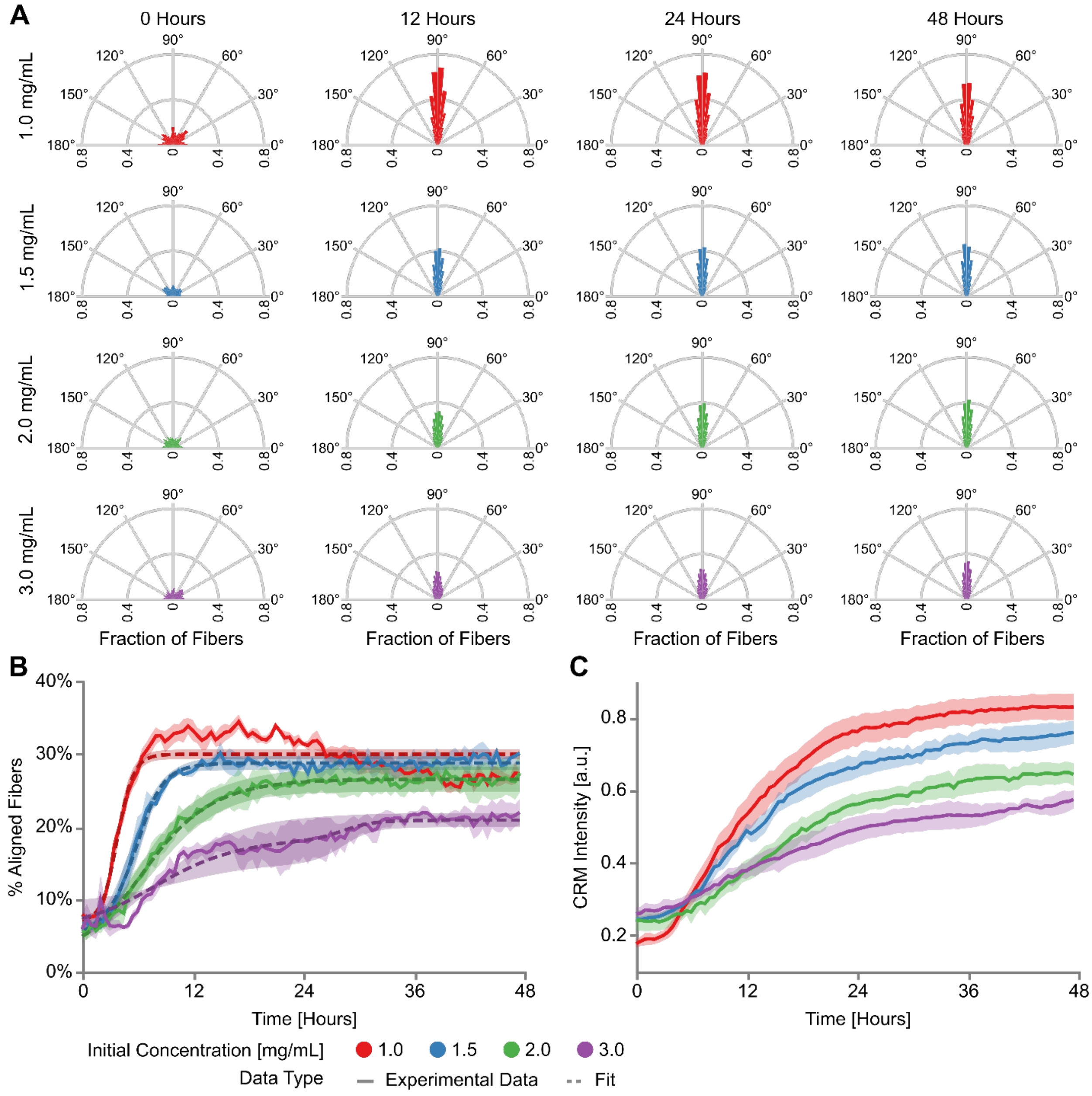


***Figure S11: Quantification of collagen microstructure from CRM*** **(A)** Polar histograms of the fraction of fiber angles quantified from confocal reflection (CRM) images using the ImageJ plugin Ridge Detection [5] on the for initial collagen concentrations 1.0, 1.5, 2.0, 3.0 mg/mL at 0, 12, 24, and 48 hours. Fiber angles between 181°-360° are wrapped around to the 0°-180° range. **(B)** Quantification of the percentage of aligned fibers (85°-95°) over 48 hours for each collagen concentration. Note the percentage of aligned fibers for the 1.0 mg/mL gels drifts downwards after

24 hours due to saturation of the CRM signal, making ridges more difficult to detect. Dashed lines show the fit logistical model as described in the "*Collagen remodeling*" section of the supplementary methods. **(C)** Average pixel intensity of CRM channel quantified as described in "*Collagen remodeling*" section of the supplementary methods.

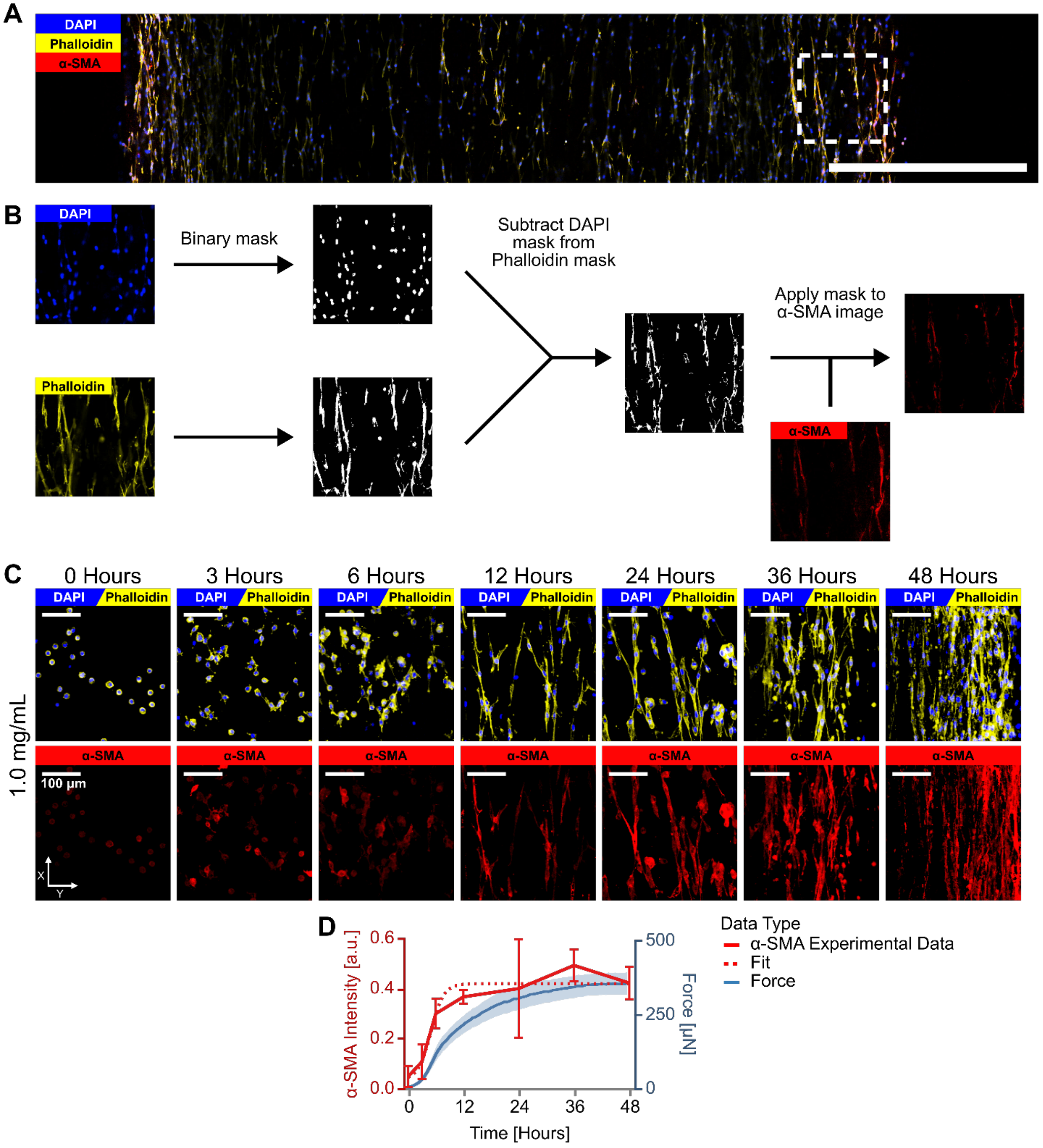


***Figure S12: Quantification pipeline of α-SMA intensity.*** (**A**) Immunofluorescence image of a uniaxially constrained gel (1.0 mg/mL, 48 hours). Blue indicates DAPI (nucleus), yellow indicates Phalloidin (F-actin), red indicates alpha smooth muscle actin (α-SMA). (**B**) Pipeline to isolate cytoplasmic α-SMA. DAPI and Phalloidin images are thresholded to generate binary masks. A cytoskeletal mask is obtained by subtracting the DAPI mask from the Phalloidin mask. The resulting cytoplasmic mask is then applied to the α-SMA image to isolate the cytoplasmic α-SMA signal. The total α-SMA intensity is calculated by summing the pixel intensity of the cytoplasmic α-SMA signal across the bottom 250 μm of the gel. Scale bar, 1000 μm. **(C)** Representative images of gels stained for DAPI and phalloidin (top row) and α-SMA (bottom row), with initial collagen concentration 1.0 mg/mL at 0, 3, 6, 12, 24, 36, and 48 hours. **(D)** Total α-SMA intensity over time is quantified using the method described in the "*Immunofluorescence*" section of the manuscript.

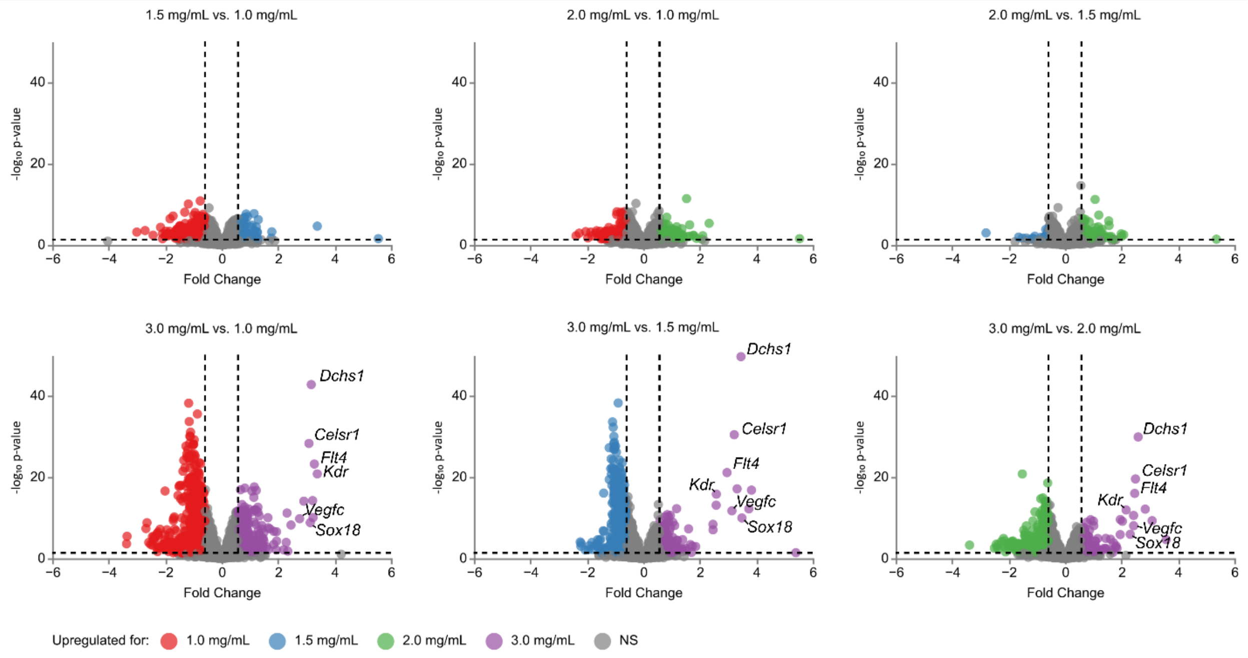


***Figure S13: RNA Seq – Volcano Plots*** Volcano plots showing differentially expressed genes (DEGs) in each pair of initial concentrations (1.0, 1.5, 2.0, 3.0 mg/mL) with $\log_2$-fold-change on the x axis and $-\log_{10}$ *p*-value on the y axis. Significant up- or downregulated DEGs (p-value < 0.05 and abs($\log_2$ Fold Change) > $\log_2(1.5)$) are shown in color and non-significant genes are shown in gray.

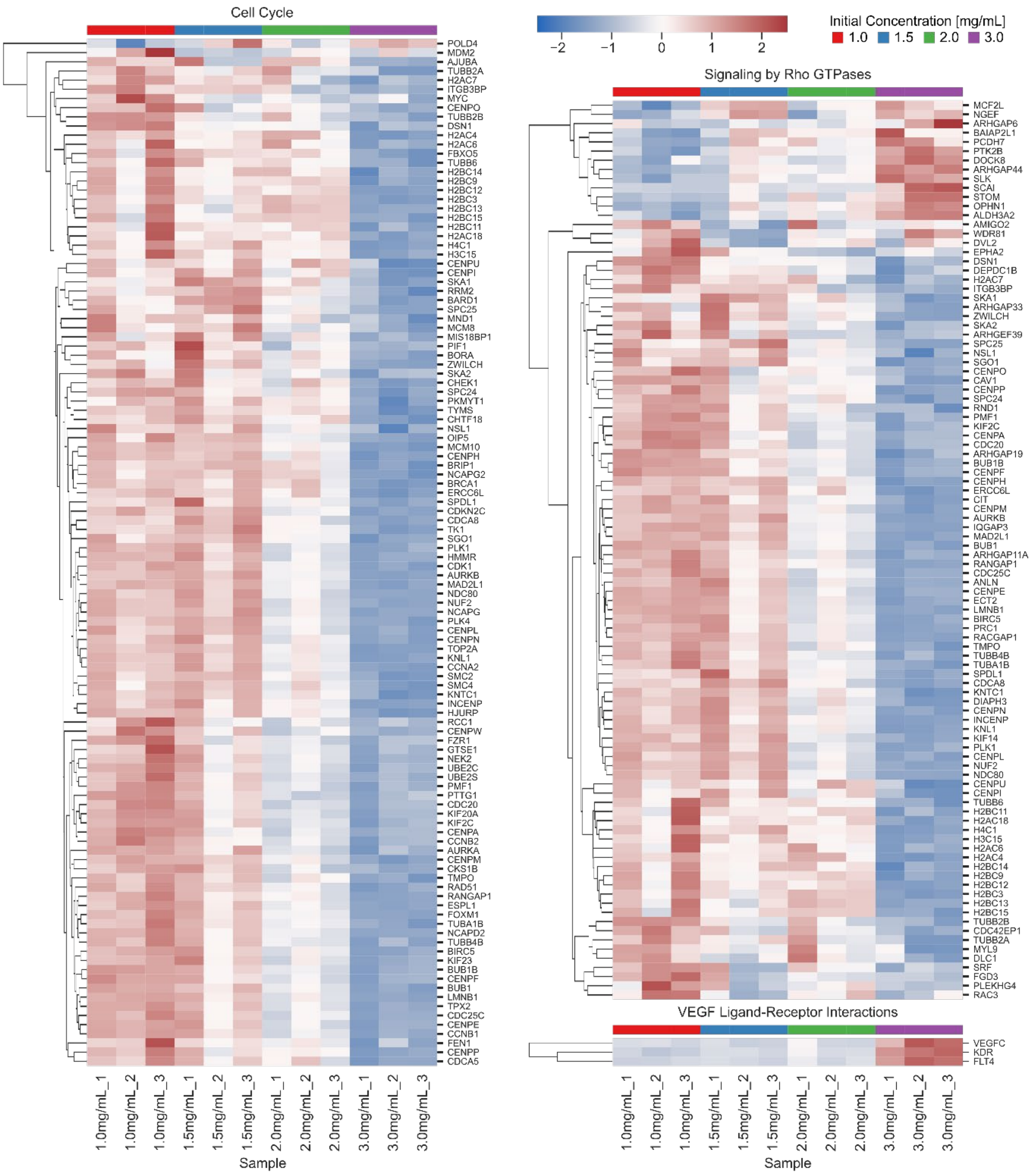


***Figure S14: RNA-seq – Heatmap*** Z-scores of normalized gene expression for the significant individual genes in the Reactome processes shown in Figure 6D.

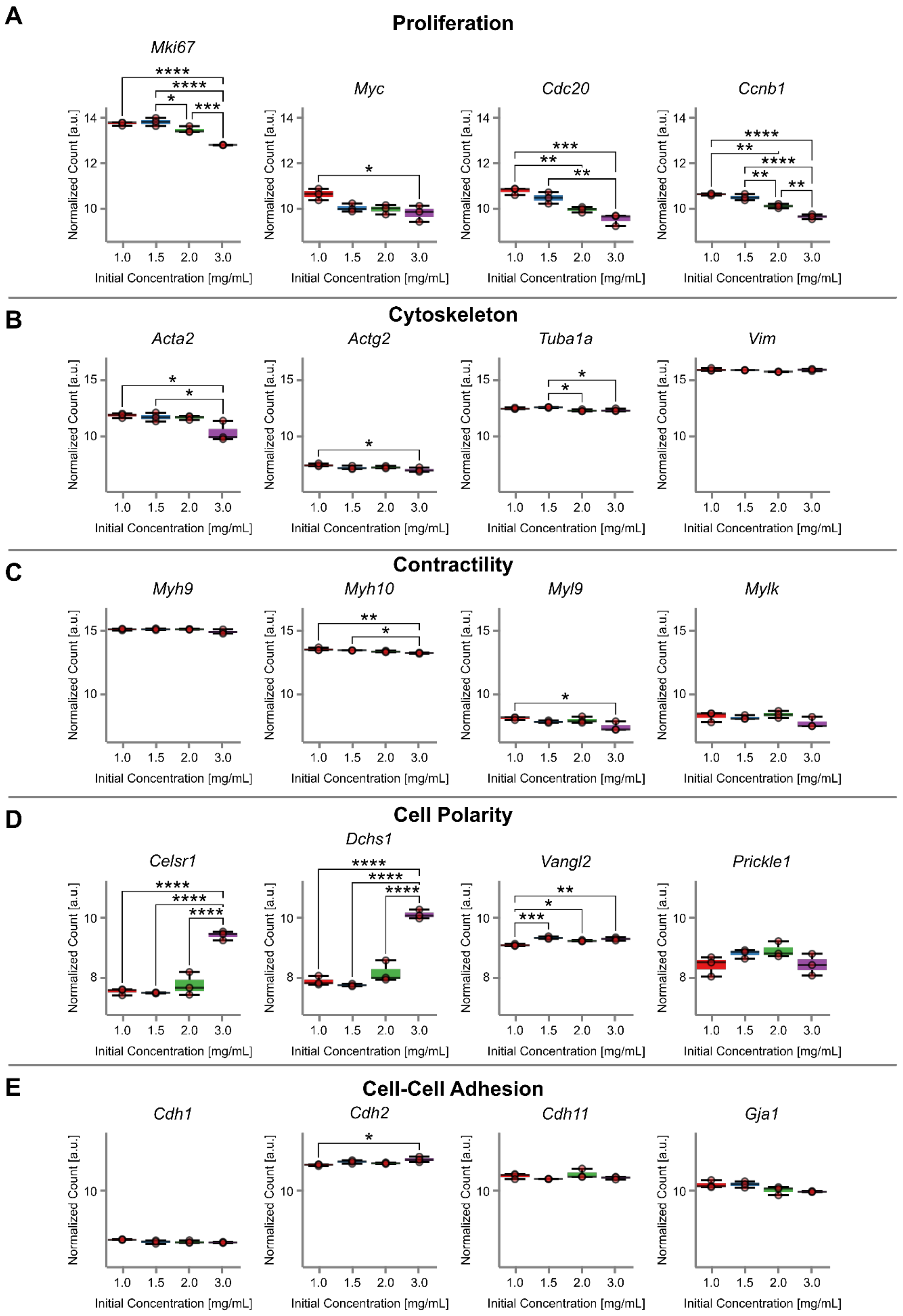
A
Proliferation
Mki67
Myc
Cdc20
Ccnb1
B
Cytoskeleton
Acta2
Actg2
Tuba1a
Vim
C
Contractility
Myh9
Myh10
Myl9
Mylk
D
Cell Polarity
Celsr1
Dchs1
Vangl2
Prickle1
E
Cell-Cell Adhesion
Cdh1
Cdh2
Cdh11
Gja1
Normalized Count [a.u.]
Initial Concentration [mg/mL]
1.0
1.5
2.0
3.0

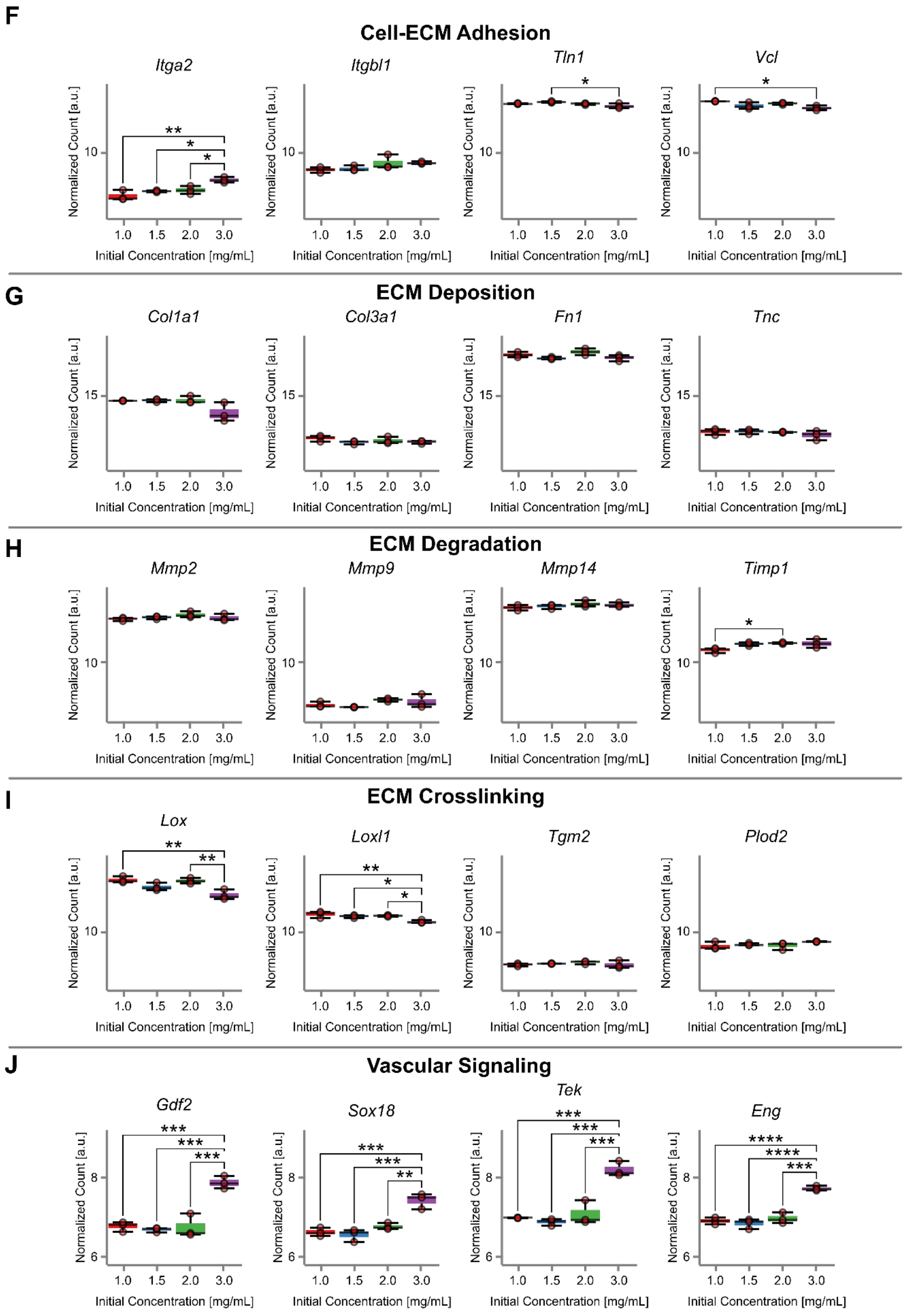
F
Cell-ECM Adhesion
Itga2
Itgbl1
Tln1
Vcl
G
ECM Deposition
Col1a1
Col3a1
Fn1
Tnc
H
ECM Degradation
Mmp2
Mmp9
Mmp14
Timp1
I
ECM Crosslinking
Lox
Loxl1
Tgm2
Plod2
J
Vascular Signaling
Gdf2
Sox18
Tek
Eng
Normalized Count [a.u.]
Initial Concentration [mg/mL]
1.0
1.5
2.0
3.0

***Figure S15: Gene expression box plots for relevant pathways*** Normalized expression of individual genes across initial collagen concentrations (1.0, 1.5, 2.0, and 3.0 mg/mL). Statistical significance was assessed using a one-way ANOVA with equal variance assumption and a Tukey post-hoc test; * $p<0.05$, ** $p<0.01$, *** $p<0.001$, **** $p<0.0001$.

### SUPPLEMENTARY VIDEOS

***Movie S1: Representative Imaging Data*** Representative 48-hour timelapse of the transmitted light detector (TD) and the Best-Z confocal reflection microscopy (CRM) images in the XY plane, as well as the CRM cross section in the YZ plane for a representative gel with 1.0 mg/mL initial collagen concentration and a cell density of 500,000 cells/mL.

***Movie S2: Cross section comparison*** Comparison of confocal reflection microscopy (CRM) cross-sections in the YZ plane for the four collagen concentrations (1.0, 1.5, 2.0, 3.0 mg/mL) over 48 hours. The segmented gel bottom is shown in red, and the arc extrapolation is shown in yellow.

***Movie S3: Representative gel retraction*** Time-lapse images capturing retraction in uniaxially constrained gels at four initial collagen concentrations (1.0, 1.5, 2.0, and 3.0 mg/mL) after 48 hours of remodeling, revealing concentration-dependent release of internal stress.

***Movie S4: Representative fiber densification and alignment*** Densification and alignment of the four collagen concentrations (1.0, 1.5, 2.0, 3.0 mg/mL) in the XY plane over a period of 48 hours.
